# Is Tau the Initial Pathology in Dopaminergic Nigrostriatal Degeneration? Studies in Parkinsonism and Parkinson’s Disease

**DOI:** 10.1101/2022.08.04.502831

**Authors:** Yaping Chu, Warren D. Hirst, Howard J. Federoff, Ashley S. Harms, A. Jon Stoessl, Jeffrey H. Kordower

**Affiliations:** ASU-Banner Neurodegenerative Disease Research Center, Arizona State University 797 East Tyler, Tempe, AZ85281, USA; Neurodegenerative Diseases Research Unit, Biogen, 115 Broadway, Cambridge, MA 02142, USA; Neurology, School of Medicine, Georgetown University Medical Center, DC 20007, USA; Department of Neurology, University of Alabama at Birmingham, AL 35294, USA; Department of Neurology, University of British Columbia BC, Canada

**Keywords:** Tau, Alpha-synuclein, Dopaminergic neurodegeneration, Parkinsonism, Parkinson’s disease

## Abstract

While Parkinson’s disease (PD) remains clinically defined by cardinal motor symptoms resulting from nigrostriatal degeneration, it is now appreciated that PD consists of multiple pathologies, but it is unclear which occurs first and which are responsible for the nigrostriatal degeneration. For the past number of years, we have been studying a well-characterized cohort of subjects with motor impairment that we have termed mild motor deficits (MMD). Motor deficits were determined on a modified and validated Unified Parkinson’s Disease Rating Scale III (UPDRS III), but they occur to a degree insufficient to diagnose PD. We consider this population to have prodromal PD. However, in past studies, cases in this cohort had a selection bias as both a clinical syndrome in between no motor deficits and PD, plus nigral Lewy pathology as defined post-mortem, were required for inclusion. Therefore, in this study, we only based inclusion on a clinical phenotype intermediate between no motor impairment and PD. Then, we divided this group further based upon whether or not they had a synucleinopathy. Here we demonstrate that loss of nigral dopaminergic neurons, loss of putamenal dopaminergic innervation, loss of TH-phenotype in the substantia nigra and putamen, and changes in axonal transport occur equally in groups with and without nigral alpha-synuclein aggregates. Indeed, the common feature of these two groups is that both have similar degrees of AT8-expressing phospho-tau, a pathology not seen in the nigrostriatal system of aged-matched controls. These finding were confirmed with early (CP13) and late (PHF1) tau markers. This suggests that the initiation of nigrostriatal dopaminergic neurodegeneration occurs independently of alpha-synuclein aggregation and is likely tau mediated.

## Introduction

Nigral dopaminergic neurodegeneration and accumulation of aggregated proteins in Lewy bodies (LB) and Lewy neurites (LN) pathologically define Parkinson’s disease (PD). The LB and LN, collectively referred to as Lewy pathology, are required for the postmortem diagnosis of definite PD[16] and are considered a precursor to neuronal degeneration[24]. According to Braak staging[5], LB deposition follows a predictable sequence, progressing in a stereotypical pattern starting caudally from the lower brainstem and moving rostrally with involvement of the substantia nigra in Braak stage 3. Alternatively, pathology may originate in the olfactory bulb. However, in Braak PD stages 1 and 2, *before* Lewy body pathology occurs in the substantia nigra, there are already reduced neuronal densities of tyrosine hydroxylase (TH) positive neurons and higher percentage of TH-immunonegative melanin-laden neurons[40]. This suggests that neurodegeneration and neuronal dysfunction precede alpha synuclein(α-syn) positive Lewy pathology in the substantia nigra. Furthermore, Lewy pathology is not always detected in the substantia nigra of PD and parkinsonian brains[6, 7, 35, 40, 58], indicating that there is an early non-synuclein pathologic process involved in PD nigrostriatal degeneration, challenging the central pathogenic role of α-syn.

Tau is a normally-occuring protein that is subject to extensive post-translational modification such as hyperphosphorylation, truncation, and glycosylation resulting in insoluble, misfolded, and aggregation of this protein. This causes disruption in the microtubule network and impairment of axonal transport, eventually causing synaptic and neuronal degeneration[41]. Tau inclusions have been found in nigral neurons by direct immunochemical studies of partially purified Lewy bodies and indirect immunohistochemical studies[33]. In some studies, 50% of PD brains have tau inclusions[59]. Further, gait impairment in older persons is associate with tau aggregation in substantia nigra[50]. PD was not initially considered to be a typical tauopathy. However, several studies have demonstrated increasing evidence of tau pathology in PD brain[2, 22, 37, 46, 59]. Genome wide association studies (GWAS) of the sporadic form of PD[43, 59] have identified *MAPT*, encoding the microtubule-associated protein tau as being associated with an increased risk of disease[51]. Whether tau is the initial pathology before Lewy pathology is an important question that to date has not been addressed.

We have studied a cohort of subjects who had minor motor deficits but a clinical syndrome insufficient for a PD diagnosis[8]. We have termed this group mild motor deficits (MMD) and believe that it represents a prodromal form of PD[8, 9, 11]. To support this concept, we have found that subjects in this cohort display decreases in TH-ir nigral neurons, decreases in putamenal TH innervation, and decreases in dopaminergic phenotype relative to subjects with no motor deficits (NMD), but not to the degree that is seen in PD[8]. However, subjects in the MMD group had 2 defining criteria; 1) a clinical motor syndrome that was measurable but insufficient to be diagnosed with PD and 2) a synucleinopathy within the substantia nigra. To eliminate this bias resulting from all cases having a synucleinopathy, we collected new cases where the only requirement was for subjects to have clinical evidence of MMD. Without bias, we assessed these cases in a blinded fashion and further categorized them as to whether a synucleinopathy was present (MMD-LB) or absent (MMD), in order to test the hypothesis that Lewy pathology was responsible for nigrostriatal degeneration. Indeed, for all our measures, MMD and MMD-LB were statistically identical. Rather strikingly, pathological tau was present in all MMD and MMD-LB cases and 22 of 24 PD cases thus may be responsible for nigrostriatal degeneration and the development of motor PD.

## Materials and methods

### Subjects

We analyzed brain tissues from older adults with no motor deficit (NMD; n=9), with mild motor deficits (n=28), and with sporadic PD (n=24). Subjects with mild motor deficits were further categorized based on the presence (MMD-LB, n=17) or absence (MMD, n=11) of nigral Lewy pathology (Table 1). Each subject signed an informed consent for clinical assessment prior to death and an anatomic gift act for donation of brain at the time of death. These subjects with NMD, MMD, and MMD-LB were participants in the Religious Orders Study, a community-based cohort study of chronic conditions of aging who agreed to brain autopsy at the time of death and were examined by a neurologist or geriatrician at the Rush Alzheimer’s Disease Center. All adults with sporadic PD were diagnosed by movement disorder specialists in the department of Neurological Sciences at Rush University Medical Center. All cases were evaluated pathologically by a board-certified neuropathologist who confirmed that these subjects did not have pathologies that met a diagnostic criteria for any other neurodegenerative disease. The Human Investigation Committee at Rush University Medical Center approved this study.

**Table 1.**
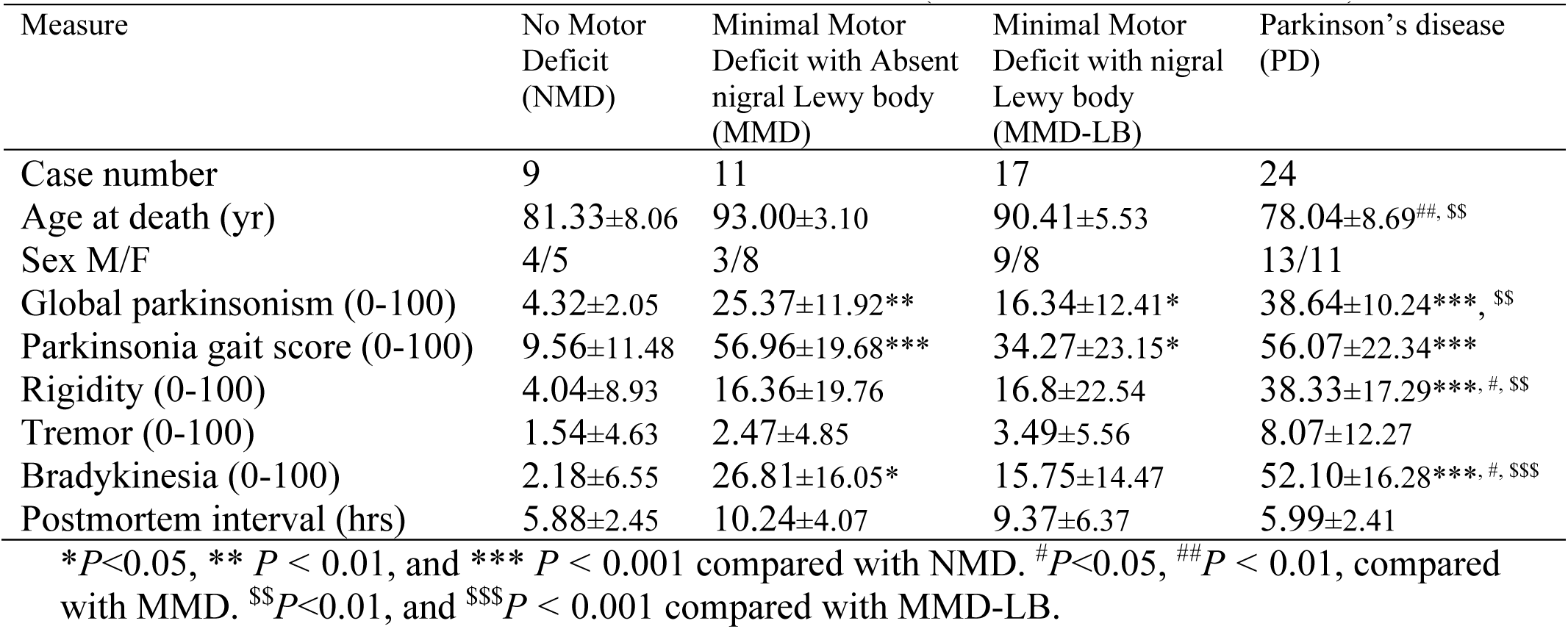
Clinical and Postmortem Characteristics (Mean ± Standard Deviation)

### Clinical evaluation and assessment of minimal motor deficits and PD

All Subjects with NMD, MMD, and MMD-LB underwent a uniform structured clinical evaluation each year that included medical history, neurologic examination, and neuropsychological performance tests[3]. Trained nurses assessed parkinsonian signs including gait, rigidity, bradykinesia, and tremor (26-items) with a modified UPDRS on a 0 to 5 scale[6], with each sign rescaled to a maximum score of 100 (see below). A summary global parkinsonian sign score (maximum possible score 100) was constructed by averaging from gait, rigidity, bradykinesia, and tremor scores (Table 1). NMD was defined as having 1 or none of these parkinsonian signs. MMD and MMD-LB were defined as having 2 or more of these parkinsonian signs with a score of 1 but with clinical features insufficient to meet the clinical definition of PD based on U.K. Brain Bank criteria. For PD, inclusion criteria included a history compatible with idiopathic PD, bradykinesia and one additional cardinal motor signs (rest tremor, rigidity, and gait disturbance). The Unified PD Rating Scale III (UPDRSIII, ON) were recorded. To compare with NMD and MMD, the 26 items included in the modified version of the UPDRS were extracted from the full UPDRSIII in patients with PD. The ratio (the raw score/the maximum possible score) was multiplied by100 (Table 1). Exclusion criteria included familial PD, the Lewy body variant of Alzheimer’s disease or the combination of PD and Alzheimer’s disease.

### Tissue processing and postmortem evaluation

At autopsy, the brains were removed from the calvarium and processed as described previously[12]. Briefly, each brain was cut into 2 cm coronal slabs and then hemisected. The slabs were fixed in 4% paraformaldehyde for 5 days at 4ºC. After 24 brain blocks were sampled from one side of the brain for pathologic diagnoses, the remaining brain slabs were cryoprotected in 0.1 M phosphate buffered saline (PBS; pH 7.4) containing 2% dimethyl sulfoxide, 10% glycerol for two days followed by 2% dimethyl sulfoxide, and 20% glycerol in PBS for at least 2 days prior to sectioning. The fixed slabs were then cut into 18 adjacent series of 40μm thick sections on a freezing sliding microtome. All sections were collected and stored at -20 ^?^C in a cryoprotectant solution prior to processing.

A complete neuropathologic evaluation was performed[50]. Dissection of diagnostic blocks included a hemisection of brain, including substantia nigra and striatum. When present, Lewy bodies were identified with H&E staining and further visualized with antibody staining for α-syn (details below) from midfrontal, midtemporal, inferior parietal, anterior cingulate, entorhinal cortex and hippocampus, amygdala, basal ganglia and midbrain. McKeith criteria[39] were modified to assess the categories of Lewy body disease. Nigral neuronal loss was estimated from midbrain. Bielschowsky silver stain was used to visualize neurofibrillary tangle in the frontal, temporal, parietal, entorhinal cortex, and the hippocampus. Braak stages were based upon the distribution and severity of neurofibrillary tangle pathology. The neuritic plaque density was scored as recommended by Consortium to establish a Registry for Alzheimer’s Disease (CERAD). In addition to the evaluation by a board-certified neuropathologist, an evaluation of Lewy pathology using α-syn immunohistochemistry allowing for the segregation of MMD into MMD and MMD-LB groups was confirmed by the lead and senior authors.

### Immunohistochemistry

An immunoperoxidase labeling method was used to visualize phosphorylated α-syn using monoclonal phospho S129 antibody (p-S129, 1:1000, ab51253, abcam), tyrosine hydroxylase (TH) expression using a monoclonal TH antibody (ImmunoStar; 1:10,000), and phosphorylated paired helical filament tau with phospho-Ser202+Thr205 using a monoclonal antibody (AT8, 1:1,000; MN1020, ThermoFisher [47]). Endogenous peroxidase was quenched by 20 min incubation in 0.1 M sodium periodate, and background staining was blocked by 1 h incubation in a solution containing 2% bovine serum albumin and 5% normal horse serum or goat serum. Tissue sections were immunostained for primary antibody overnight at room temperature. After 6 washes, sections were sequentially incubated for 1 h in biotinylated horse anti-mouse IgG (for TH and AT8) or goat anti-rabbit IgG (for p-S129) followed by the *Elite* avidin–biotin complex (1:500; Vector) for 75 min. The immunohistochemical reaction was completed with 0.05% 3, 3′-diaminobenzidine (DAB) and 0.005% H2O2. Stained sections were mounted on gelatin-coated slides, dehydrated through graded alcohol, cleared in xylene, and coverslipped with Cytoseal (Richard-Allan Scientific, Kalamazoo, MI).

### Evaluating densities of TH immunoreactive neurons and AT8 and p-S129 immunoreactive aggregates in substantia nigra

The density of nigral TH-immunoreactive (TH-ir) neurons and misfolded proteins were estimated for each subject. All stereological estimates were separately counted using a uniform, systematic, and random design. An optical fractionator unbiased sampling design was used and a Cavalieri’s principle to assess the volume within substantia nigra[10, 13, 27]. In each subject, we evaluated the substantia nigra pars compacta from the level of the midbrain at the exit of the 3rd nerve to the decussation of the superior cerebellar peduncle. Approximately 5 equispaced sections were sampled from each brain. The section sampling fraction (ssf) was 1/0.055. The distance between sections was approximately 0.72 mm. In cross-section, the substantia nigra is in the ventral midbrain. The substantia nigra pars compacta was outlined using a 1.25× objective. A systematic sample of the area occupied by the substantia nigra pars compacta was made from a random starting point (StereoInvestigator v2021.1.3 software; Micro-BrightField, Colchester, VT). Counts were made at regular predetermined intervals (x = 313μm, y = 313μm), and a counting frame (70 × 70μm = 4900 μm^2^) was superimposed on images obtained from tissue sections. The area sampling fraction (asf) was 1/0.05. These sections were then analyzed using a 60× Planapo oil immersion objective with a 1.4 numerical aperture. The section thickness was empirically determined. Briefly, as the top of the section was first brought into focus, the stage was zeroed at the z-axis by software. The stage then stepped through the z-axis until the bottom of the section was in focus. Section thickness averaged 16.21 ± 2.3μm in the midbrain. The disector height (counting frame thickness) was 10 μm. This method allowed for 1μm top guard zones and at least 2μm bottom guard zones. The thickness sampling fraction (tsf) was 1/0.62. Care was taken to ensure that the top and bottom forbidden planes were never included in the cell counting. The ultimate estimate of the counted profiles within the substantia nigra pars compacta was calculated separately using the following formula: N=ΣQ−·1/ssf · 1/asf · 1/tsf. ΣQ− was the estimated number of raw counts.

The Cavalieri estimator (StereoInvestigator software; Micro-BrightField VT) was used to estimate volume of the substantia nigra pars compacta on immunostained sections. We sampled serial sections of substantia nigra that extended from the exit of the 3rd nerve to the decussation of the superior cerebellar peduncle using the optical fractionator principle described above. The distance between sections interspace was approximately 0.72 mm. According to NM the substantia nigra is outlined in cross-section using a 1.25x objective. The section thickness was empirically determined. Section thickness averaged 16.23±2.2μm. The area estimation of substantia nigra pars compacta was performed by means of a 50 × 50 μm point grid with 10x objective. The total volume of substantia nigra pars compacta was calculated by Cavalieri estimator software[12, 27]. The coefficients of error (CE) were calculated according to the procedure of Gunderson and colleagues as estimates of precision [49, 56]. The values of CE were 0.12 ± 0.05 (range 0.10 to 0.15) in PD and 0.10 ± 0.02 (range 0.08 to 0.12) in MMD and NMD.

### Evaluating densities of TH and AT8 immunoreactivities in putamen

Quantification of the relative optical density of putamenal TH immunoreactivity was performed using densitometry software (ImageJ; National Institutes of Health, Bethesda, MD), as described previously[8]. Using stereological principles, the putamen was outlined with 1.25x objective on 5 stained sections through rostral to caudal putamen and more than 100 images were randomly sampled with 20x objective for each subject. Optical density measurements for TH immunoreactivities were performed in greyscale (0 represented a maximum bright image and 255 represented a maximum dark image). To account for differences in background staining intensity, background optical density measurements in each section were taken from corpus callosum, which lacked a TH-positive profile. The mean of these measurements constituted the background optical density that was subtracted from the optical density of TH-immunoreactive intensity measurements to provide a final optical density value.

Measurements for AT8-ir punctuated dots and threads were performed using particle analyses program of Image J[32]. The black and white images were taken from AT8-stained putamenal sections using stereological principle with 20x objective. The scale was 2.64 pixels/micron. All images were thresholded at default and analyzed particles at 1 pixel size or above. The densities of AT8-ir particles were calibrated in square millimeter (mm^2^) of putamen.

### Double and triple staining and evaluating optical densities of immunofluorescence

A double-label immunofluorescence procedure was employed to determine whether tau aggregates co-existed in neurons that expressed α-syn and whether tau aggregates affected the dopaminergic neuronal functions. Midbrain and putamenal sections were incubated in the first primary antibody AT8 overnight and the goat anti-mouse antibody coupled to DyLight 649 (1:200, Jackson ImmunoResearch) for 1 h. After blockade for 1 h, the sections were then incubated in the second primary antibodies (p-S129, 1:500; TH, P40101/Pel-Freez, 1:1000; kinesin light chain (KLC) AP8637c/ABGENT, 1:500) overnight, and the goat anti-rabbit antibody coupled to DyLight 488 (1:200) for 1 h. The stained sections were mounted on gelatin-coated slides, dehydrated through graded alcohol, cleared in xylene, and covered using DPX (Sigma-Aldrich). A triple-label immunofluorescence procedure was employed to examine three and four repeat (3R, 4R) tau isoforms and their colocalization with AT8. Midbrain sections were incubated overnight in primary antibodies: 3R-tau (rat monoclonal, 1:500, 016-26561/ Wako), 4R-tau (rabbit monoclonal, 1:500, ab218314/abcam), and AT8 (mouse monoclonal, 1:1000) and for 2 hours in secondary antibodies: Cy2 conjugated goat-anti rat (1:200), Cy3 conjugated goat-anti rabbit (1:200), and Cy5 conjugated goat-anti mouse (1:200). The stained sections were mounted on gelatin-coated slides, dehydrated through graded alcohol, cleared in xylene, and covered using DPX (Sigma-Aldrich). Fluorescence intensity measurements were performed according to our previously published procedures [9, 13]. All immunofluorescence labeled images were scanned with an Olympus Confocal Fluoroview microscope equipped with argon, helium-neon lasers, and transparent optics. At 20× magnification objective and a 488, 543, and 633 nm excitation source, images were acquired at each sampling site in the substantia nigra pars compacta and were saved to a Fluoroview file. Following acquisition of an image, the stage moves to the next sampling site to ensure a completely non-redundant evaluation. Once all images were acquired, optical density measurements were performed on individual nigral neurons. To maintain consistency of the scanned image for each slide, the laser intensity, confocal aperture, photomultiplier voltage, offset, electronic gain, scan speed, image size, filter and zoom were set for the background level whereby autofluorescence was not visible with a control section. These settings were maintained throughout the entire experiment [13]. The intensity mapping sliders ranged from 0 to 4095; 0 represented a maximum black image and 4095 represented a maximum bright image. The TH-ir or KLC-ir perikarya with or without AT8-ir aggregates were identified and outlined separately. Quantitative optical density of immunofluorescence was performed on individual TH-ir or KLC-ir soma with or without AT8-ir aggregates in different channels. Five equispaced sections of the substantia nigra were sampled and evaluated. The number of cells per case was analyzed as follows: 50–70 nigral cells that contained AT8-ir aggregates and >100 nigral cells that did not contain aggregates per subject. To account for differences in background staining intensity, five background intensity measurements lacking immunofluorescent profiles were taken from each section. The mean of these five measurements constituted the background intensity that was then subtracted from the measured optical density of each individual neuron to provide a final optical density value. To confirm co-localization of the AT8 and p-s129-α-syn immunofluorescence, optical scanning through the neuron’s z-axis was performed at 1-μm thickness and neurons suspected of being double labeled were confirmed with confocal cross-sectional analyses. The optical density of TH-immunofluorescent intensities in putamen was determined using Image J program described above.

### Data analyses

Clinical characteristics were compared across groups using a Kruskal-Wallis ANOVA and where appropriate by followed by a Dunn’s Multiple post-hoc test. neuronal number, aggregates number, particle densities, and optical density measurements were compared across groups using one-way ANOVA followed by Tukey post hoc tests controlling for multiple comparisons (Prism 4, GraphPad Software, Inc.). The level of significance was set at 0.05 (two-tailed).

### Digital illustrations

Conventional light microscopic images were acquired using an Olympus microscope (BX61) attached to a Nikon digital camera DXM1200 and stored as tif files. Confocal images were exported from the Olympus laser-scanning microscope with Fluoview software and stored as tif files. All figures were prepared using Photoshop 7.0 graphics software. Only minor adjustments of brightness were made.

## Results

### Parkinsonian signs in subjects with and without a clinical diagnosis of PD

We analyzed brain tissues from nine older adults with NMD, eleven older adults with MMD, seventeen older adults with MMD-LB, and twenty-four adults with a clinical diagnosis of PD. The diagnoses of subjects with MMD, MMD-LB, and PD were confirmed pathologically in each case and there was no evidence of atypical Parkinsonism (e.g. PSP, CBD, or MSA) in any subject. Demographics are provided in Table 1. NMD subjects displayed low global scores as well as low scores on individual signs of gait, bradykinesia, rigidity, and tremor. In contrast, subjects with PD exhibited significantly higher scores on each of these measures. For the MMD and MMD-LB cases, the bradykinesia and parkinsonian gait were significantly higher as compared with NMD. Other scores were intermediate but no statistical differences between groups were observed. There was no difference on tremor score among groups. Detailed clinical examinations from the individuals are presented in supplementary Table1.

### Qualitative and quantitative observations of nigral phosphorylated *α*-syn immunoreactivity

To verify that subjects from MMD group had no nigral Lewy pathology diagnosed by the neuropathologists, we examined the phosphorylated α-syn labelling in substantia nigra in all of participants. Qualitative and quantitative analyses revealed that phosphorylated α-syn aggregates were widely distributed through substantia nigra in MMD-LB (355.59±210.01/mm^3^) and PD (398.69±227.15/mm^3^) groups but not in NMD and MMD groups (supplementary Fig. 1). These data further demonstrate that MMD subjects with motor deficits did not display nigral α-syn pathology.

### Characteristics of tyrosine hydroxylase expression in nigrostriatal system of groups

*SUBSTANTIA NIGRA*. Subjects from NMD group had a high density of TH-ir soma and an intricate local plexus of TH-ir processes within the substantia nigra (Fig. 1A, 1B). A few NM-laden nigral neurons were TH-immunonegative. MMD subjects showed a clear reduction in TH-ir neurons (Fig. 1E). Many NM-laden nigral neurons were TH-immunonegative and the TH-ir neurons displayed less extensive processes (Fig 1F). MMD-LB subjects also displayed obvious reductions of TH-immunoreactivity and qualitatively this reduction appeared identical to the MMD cases. (Fig. 1I). As with the MMD cases, many NM-laden neurons in the MMD-LB cases displayed TH-immunonegative perikarya and decreased TH-ir arborization. (Fig. 1J). In PD subjects, both TH-ir somata and dendrites in substantia nigra (Fig. 1M, 1N) were severely reduced to a degree greater than what was seen in subjects with MMD and MMD-LB.

**Figure 1.**
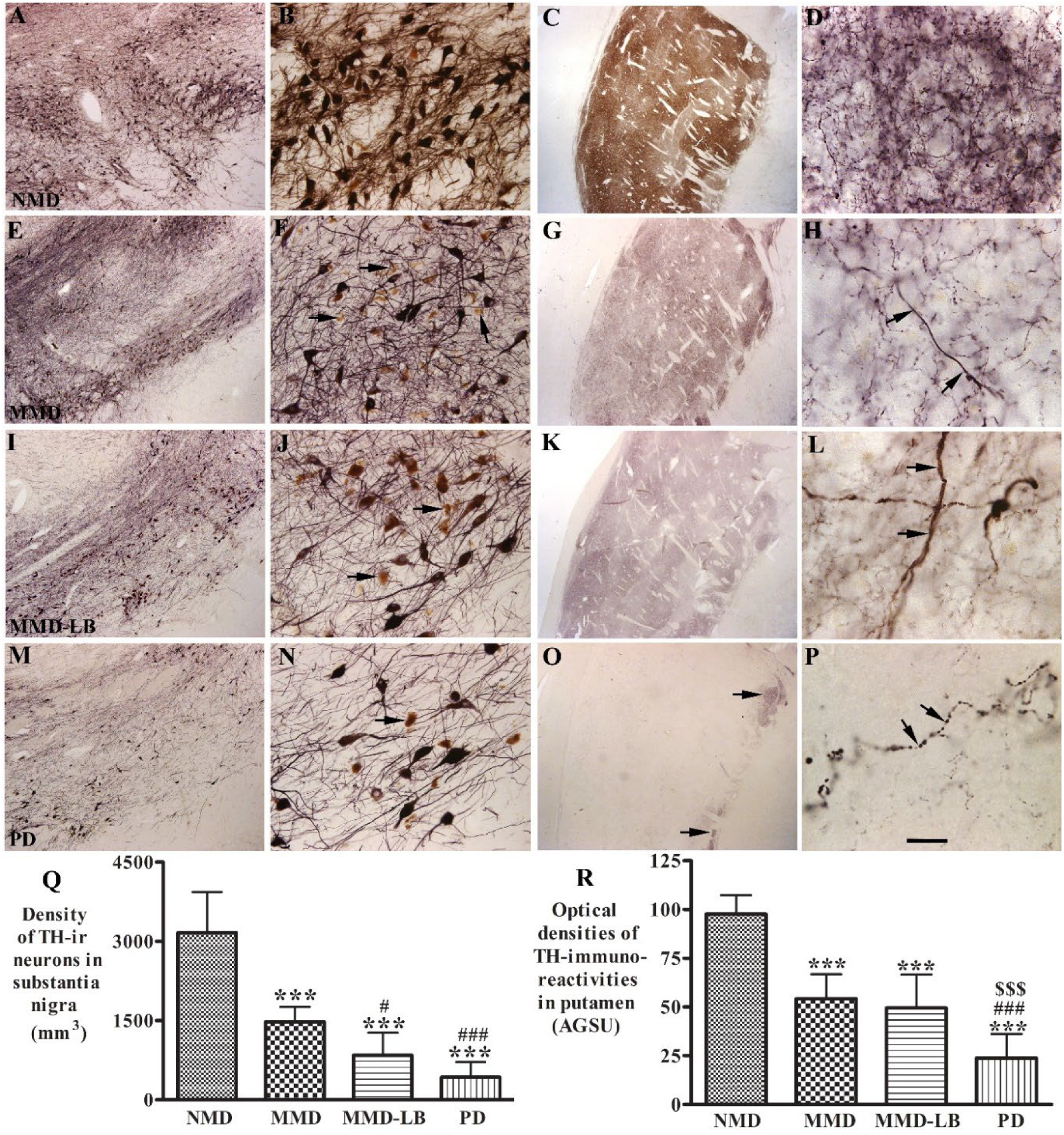
Qualitative and quantitative evaluation for nigral and putamenal TH expression. Photomicrographs of substantia nigra (left two columns) and putamen (right two columns) from no motor deficit (NMD; A-D), minimal motor deficits (MMD; E-H), minimal motor deficits with Lewy body (MMD-LB; EI-L), and Parkinson’s disease (PD; M-P) illustrate TH immunoreactivity. Subject from the NMD group showed intense TH-immunoreactive (TH-ir) somata with extensive local plexus of TH-ir processes (A, B) in substantia nigra and dense TH-ir fiber throughout putamen (C, D). Nigral TH immunoreactivities were reduced in MMD (E, F) and MMD-LB subjects (I, J) compared with NMD (A, B) and some remaining nigral melanized neurons exhibited no detectable TH immunoreactivity (arrows; F, J). Putamenal TH immunoreactivities were reduced in MMD (G, H) and MMD-LB (K, L) and some of remaining TH-ir fibres displayed swollen varicosities (arrows; H, L). PD cases displayed severe reduction of TH immunoreactivity in substantia nigra (M. N) and undetectable TH immunoreactivity in major putamen, except the ventromedial putamen near globus pallidus (arrows; O) and few remaining TH immunoreactive fibres exhibited swollen segments (arrows; P). Scale bar = 20 μm in P (applies to D, H, L); 2.0 mm for C, G, K, O; 100 μm for B, F, J, N; 500 μm for A, E, I, M. (Q) Stereological analyses revealed that the density of TH-positive neurons was gradually reduced from MMD, MMD-LB to PD relative to NMD group. \*\*\**p<* 0.001, compared with NMD. ^#^*p*<0.05 ^###^*p*<0.001 compared with MMD. Densitometry (R) demonstrated that the TH-immunoreactive intensity in putamen was accordingly reduced from MMD, MMD-LB to PD relative to NMD group. \*\*\**p<* 0.001, compared with NMD. ^###^*p*<0.001 compared with MMD; ^$$$^*p*<0.001 compared with MMD-LB.

Stereological analyses confirmed that the densities of TH-ir neurons were decreased in PD cases (430.14±283.91/mm^3^) relative to and NMD subjects (3168.48±770.76/mm^3^). There were similar, but smaller reductions in TH-ir nigral neurons in the MMD (1479.38±282.70/mm^3^) and MMD-LB (842.53±435.76/mm^3^) relative to the NMD group. Relative to the NMD group, the reduction of nigral TH-ir neurons was 53.30% in MMD, 73.29% in MMD-LB, and 85.78% in PD. An ANOVA test revealed a statistically significant difference across these groups (Fig. 1Q; p<0.0001). Post hoc analyses demonstrated a significant reduction of TH-labeled neurons with MMD (P<0.001) MMD-LB (P<0.001) and PD (p<0.001) compared with NMD group, and between MMD and PD groups (P<0.001). Critically, there was no significant difference between MMD and MMD-LB (P>0.05) groups indicating that Lewy pathology does not affect the loss of TH-ir neurons (in MMD) and suggests that factors others from Lewy pathology may be responsible for nigral cell loss.

#### TH-innervation of the putamen

TH-labeled dopaminergic terminals from NMD subjects were distributed in a mosaic pattern of low- and high TH-ir zones (Fig. 1C). Dense fine TH-ir fibers were distributed throughout the grey matter of putamen, consisting of a fine mesh of fiber (Fig. 1D). In contrast, TH-labelling from MMD subjects (Fig. 1G, 1H) was remarkably decreased compared with NMD controls (Fig. 1C, 1D). The TH-ir fine fibers were obviously reduced (Fig. 1H) although the light TH-labeling mosaic pattern remained (Fig. 1G). The TH-ir pattern from MMD-LB (Fig. 1K, 1L) and MMD (Fig. 1G) appeared similar. In both MMD groups, TH-ir fine fibers were diminished relative to NMD subjects and remaining putamenal fibers often displayed an abnormal morphology characterized by swollen varicosities (Fig. 1H, 1L). In each of the PD cases, TH-ir fine fibers were barely detectable in the putamen (Fig. 1O, 1P) and the few remaining thick fibers displayed swollen varicosities and varicose segments (Fig. 1P). A few fine TH-ir fibers were observed in the more ventromedial putamen near globus pallidus (Fig. 1O).

Optical densities supported the qualitative observations revealing that relative to the NMD group, putamenal TH optical densities were reduced 44.41% in MMD, 49.26% in MMD-LB, and 75.59% in PD group. An ANOVA revealed a statistically significant difference across these groups (Fig. 1R; p<0.0001). Post hoc analyses demonstrated a significant reduction of putamenal TH-ir intensities in subjects with MMD (P<0.001), MMD-LB (P<0.001), and PD (p<0.001) compared with NMD group, between MMD and PD group (P<0.001), and between MMD-LB and PD (p<0.001). Critically, there was no difference between the MMD and MMD-LB groups (P>0.05) supporting the concept that α-syn pathology is not necessary for the loss of TH-innervation and suggests that factors other than Lewy pathology are responsible for putamenal loss of dopaminergic innervation.

### Morphological characteristics of phosphorylated tau invading into substantia nigra and putamen

Since α-syn pathology does not appear to be necessary for nigrostriatal degeneration in subjects with MMD, we examined the potential for tau to fill this role, since tau has been shown previously to be a co-pathology in PD[45, 46], and knockout of tau attenuates, and delays neurodegeneration associated with α-syn [53]. We examined tau pathology in all participants in this study. Phosphorylated tau aggregates were observed in the nigrostriatal system in all MMD and MMD-LB subjects and 22 of 24 PD subjects with PD. Immunohistochemistry with AT8 antibody revealed multiple patterns of AT8-ir aggregates within the substantia nigra. Some nigral neurons had small AT8-labeled punctate granules (Fig. 2A, 2B) in highly melanized neurons. The distribution of neuromelanin (NM) in cells with and without AT8-labeled punctate granules appeared similar. As phosphorylated tau accumulates, AT8-labeled clumpy granules fill the perikarya, (Fig. 2C). Under certain circumstances, the phosphorylated tau filled the whole cell including perikarya and proximal processes (Fig. 2D) and the NM was not visible. Finally, the phosphorylated tau was seen to be compressed into a spherical shape (Fig. 2E). AT8-labeled processes displayed swollen threads and segments distributed through the substantia nigra (Fig. 2F).

**Figure 2.**
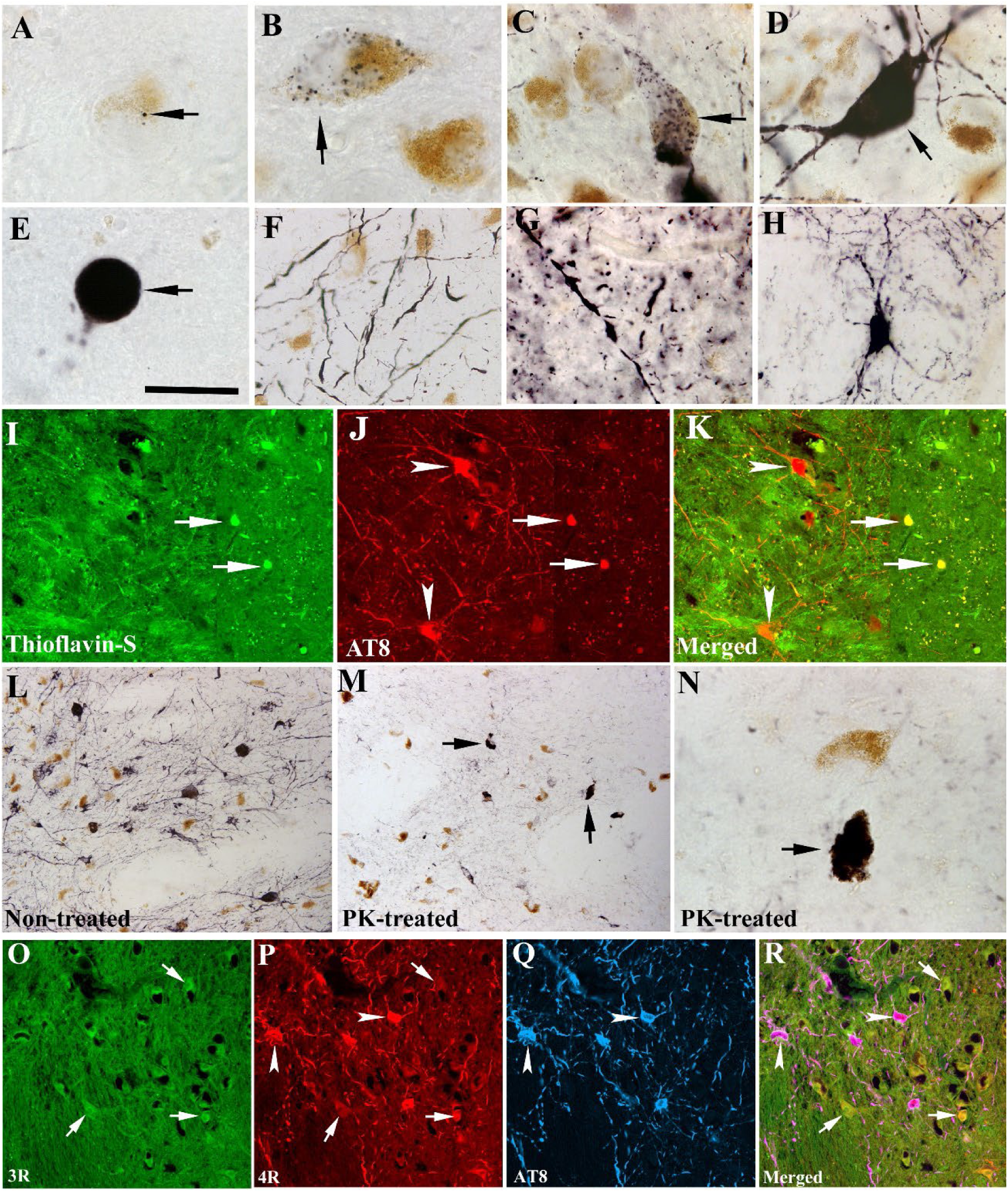
Morphologic features of tau aggregation in nigrostriatal system. Photomicrographs of substantia nigra (A-F) and putamen (G, H) show the shapes of AT8-immunoreactive (AT8-ir) aggregates. AT8-ir punctate granule was seeded (black; arrow; A) and mixed (black; arrow; B) into neuromelanin (brown). AT8-ir clumpy granules were accumulated within perikarya (arrows; C) or filled with whole neuron including somata and main processes (arrow; D) that the NM was invisible. AT8-ir sphere aggregate like a Lewy body (arrow; E) with cytoplasm and processes disappear. AT8-ir fragmental threads (F) were distributed in substantia nigra. AT8-ir varicosities and punctuated boutons dispersed into putamen (G). Few AT8-ir putamenal neuron displayed dark stained somata with abundant processes (H). Double-labelling revealed the dense phosphorylated tau accumulations (AT8, red; arrows; J) were thioflavin-S positive (Green, arrows; I) but the granules phosphorylated tau were thioflavin negative (arrowheads, I-K). Phosphorylated tau immunolabeling (L) observed from the section without proteinase K (PK) treatment was basically eliminated following PK treated section (M, N). However, the dense tau accumulations (arrows; M, N) were resistant to PK treatment. Fluorescent triple-labelling (O-R) showed that both 3R (green, arrows; O) and 4R (red, arrows and arrowheads; P) tau isoforms existed in nigral melanized neurons. 4R isoform in same melanized neurons displayed stronger intense staining than 3R and colocalized with phosphorylated tau marker AT8 (blue, arrowheads; P-R). Scale bar in E =20 μm (applies to A-H, N), 100 μm in I-K, and 120 μm in L, M, O-R.

In the putamen, AT8-labeled products exhibited swollen varicosities, segments, and punctate boutons (Fig. 2G). The swollen varicosities present as a string of beads-like axonal swelling. The punctate boutons appeared as granules distributed into putamenal grey matter. Fewer AT-8 labelled cells (Fig. 2H) displayed neuronal perikarya with oval or triangular shape and extensive local processes.

Double staining revealed that the dense tau aggregates were oftebn thioflavin S-positive, but dispersed tau granules in perikarya and processes were not (Fig. 2I-2K). Proteinase K digestion was used to further determine whether the AT8 labeled tau was soluble (non-aggregated) or insoluble (inclusions). In this experiment, proteinase K digested the granular tau structures but not the dense tau accumulation (Fig. 2L-2N). These observations indicate that tau pathology progresses in nigrostriatal system during prodromal PD with apparent transition into aggregates that are resistant to protein K digestion.

The adult human brain expresses multiple main tau isoforms, which can be categorized as 3R or 4R tau based on whether they contain three or four carboxy-terminal repeat domains[38]. 3R and 4R tau is more common in Alzheimer’s disease while 4R Tau is seen in Progressive Supranuclear Palsy[19]. To determine which isoform exists in the substantia nigra, the 3R and 4R isoforms were examined and analyzed with colocalization of AT8-labeled phosphorylated tau. Fluorescent labelling revealed that both 3R (Fig. 2O) and 4R (Fig. 2P) tau isoforms were distributed in nigral NM-laden neurons.

### Qualitative and quantitative observations of phosphorylated tau in nigrostriatal system

Our previous study[8] indicated that the density of phosphorylated α-syn aggregates in subjects with MMD-LB was similar to PD. We now examined the distribution of tau inclusions from subjects with minimal motor deficits and clinically diagnosed PD. In this regard, AT8-labeled aggregates in substantia nigra and neuropil threads in putamen were examined and quantified in all subjects.

The density of AT8-labeled aggregates was variable from case to case. Stereological analyses showed that the density of nigral AT8-labled aggregates was higher in subjects with MMD (223.78±169.61/mm^3^) and MMD-LB (228.4±196.57/mm^3^), medium in the PD (151.52±165.75/mm^3^), and much lower in NMD (8.07±17.54/mm^3^). An ANOVA revealed a statistically significant difference across these groups (Fig. 3Q; p<0.001). Post hoc analyses demonstrated a significant higher density of AT8-labeled aggregates in MMD (P<0.001), MMD-LB (P<0.001) and PD (p<0.001) compared with NMD group, between MMD and PD (P<0.001) between MMD-LB and PD (P<0.001), but there was no difference between MMD and MMD-LB groups (P>0.05).

**Figure 3.**
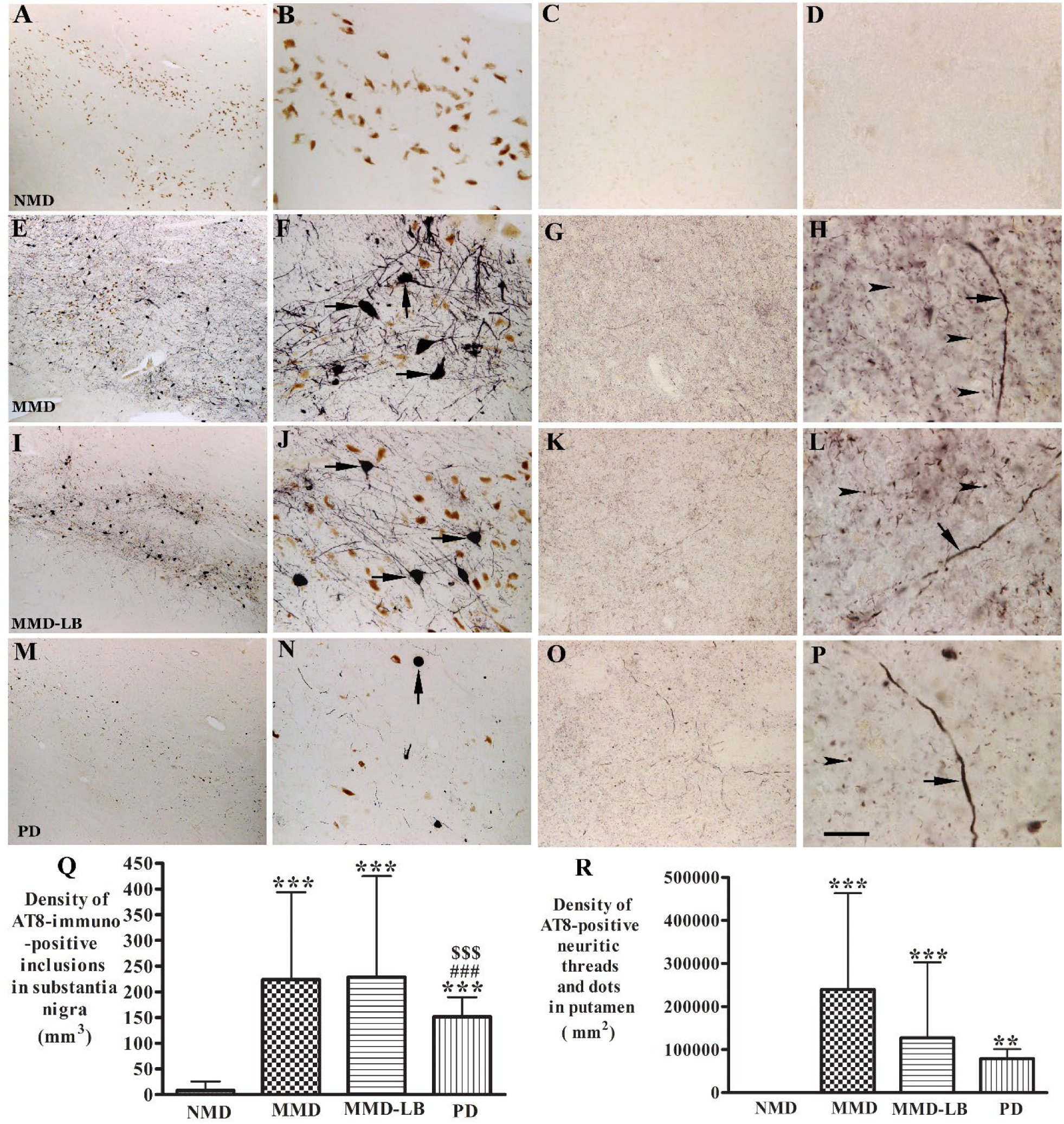
Qualitative and quantitative evaluation for tau aggregates in substantia nigra and putamen. Photomicrographs of the mid-substantia nigra (left two columns) and putamen (right two columns) from no motor deficit (NMD; A-D), minimal motor deficits (MMD; E-H), minimal motor deficits with nigral Lewy body (MMD-LB; I-L), and Parkinson’s disease (PD; M-P) show AT8-immunoreactive (AT8-ir) patterns. AT8 immunoreactivity was not detected in NMD group (A-D). In contrast, numerous AT8-ir neurons were distributed throughout substantia nigra (E, I) and displayed dark somata with abundant processes in subjects with MMD (arrows; F) and MMD-LB (arrows; J). AT8-ir intensities in putamen were higher in MMD (G) and MMD-LB (K) and displayed punctuated dots (arrowheads; H, L) and same expanded fiber (arrow; H, L). The AT8-ir aggregates with limited processes (arrow; N) in substantia nigra (M) and relative lesser AT8-ir punctuated dots (arrowhead; P) in putamen were observed in PD. Scale bar in P = 20 μm (applies to D, H, L); 100 μm for B, C, F, G, J, K, N, O; 500 μm for A, E, I, M. (Q) Stereological analyses revealed that the density of AT8-ir aggregates in substantia nigra and (R) the density of AT8-ir dots and threads in putamen were significant higher in MMD, MMD-LB, and PD relative to NMD group. \*\**p<* 0.01, and \*\*\**p <* 0.001 compared to NMD, ^###^*p*<0.001 compared to MMD, and ^$$$^ *p*<0.001compared to MMD-LB.

Tau phosphorylation was further investigated using two others commonly employed phosphorylated-tau antibodies, CP13 (phospho-serine 202) and PHF-1 (phosphor-serine396/serine404). Immunohistochemistry revealed that both CP13 and PHF-1labeled tau were widely distributed in substantia nigra in subjects with MMD, MMD-LB, and PD but not in the NMD subjects (Supplementary Fig.2 and Fig.3). There were more CP13, and PHF-1 labeled aggregates in subjects with MMD and MMD-LB than PD. The pattern of CP13 and PHF-1labeled tau in substantial nigra is similar to AT8 staining indicating that multiport phosphorylation sites contribute to tau aggregation in nigral dopaminergic neurodegeneration.

Within the putamen, AT8 immunoreactivity was virtually undetectable in NMD subjects (Fig. 3C, 3D). Different intensities of AT8 immunoreactivities were observed in putamen of each subject with MMD, MMD-LB, or PD. Both MMD and MMD-LB subjects had intense AT8-labeled products that were characterized by densely puncta, boutons, segments, and swollen varicosities (Fig. 3G, 3H, 3K, 3L). The density of AT8-labeled punctuated boutons in PD group (Fig. 3O, 3P) was relatively lower than seen in the MMD and MMD-LB groups (Fig. 3H, 3L). Quantitative analyses showed that the density of nigral AT8-labled aggregates was highest in subjects with MMD (239891.2±223426.5/mm^2^), higher in MMD-LB (127876.3±174805.8/mm^2^), medium in the PD (78948.9±99519.2/mm^2^), much lower in NMD (40.37±114.19/mm^2^). An ANOVA revealed a statistically significant difference across these groups (Fig. 3R; P<0.0001). Post hoc analyses demonstrated a significantly higher density of AT8-labeling in MMD (P<0.001), MMD-LB (P<0.001), and PD (p<0.01) compared with NMD group but there was no difference among MMD, MMD-LB, and PD groups (P>0.05). These data indicate that tau pathology is present in the development of nigrostriatal degeneration, especially during the early stages and in cases in which Lewy pathology does not exist.

### Co-localization analysis of tau and *α*-synuclein inclusion in the nigrostriatal system

Findings of tau accumulation in nigrostriatal system of subjects with MMD-LB and PD above led us to hypothesize that there was a co-occurrence of tau and α-syn aggregates in dopaminergic neurodegeneration in these populations. In this regard, both tau and α-syn aggregations were examined using double-labeling with AT8 and p-S129-α-syn antibodies. Co-localization analyses demonstrated three neuronal populations in the substantia nigra: 1) neurons that co-localize AT8 and p-S129-α-syn (Fig. 4A-C); 2) neurons with AT8-labeled inclusions alone (Fig. 4B, 4C), or 3) p-S129-α-syn labeled aggregates alone (Fig. 4A, 4C). Cross-sectional analyses of co-localization further revealed that the p-S129-α-syn inclusion deposited into neurons with AT8-staining somata and processes (Fig. 4D-G). However, only 5-8% of AT8-labeled aggregates co-localized with p-S129-α-syn within nigral neurons. The majority of AT8 staining was not co-localized with α-syn. In putamen, longer neuropil threads staining for AT8 were p-S129 immunopositive (Fig. 4H-J) but short segments and puncta were not. These data are consistent with the notion that tau is driving nigrostriatal degeneration.

**Figure 4.**
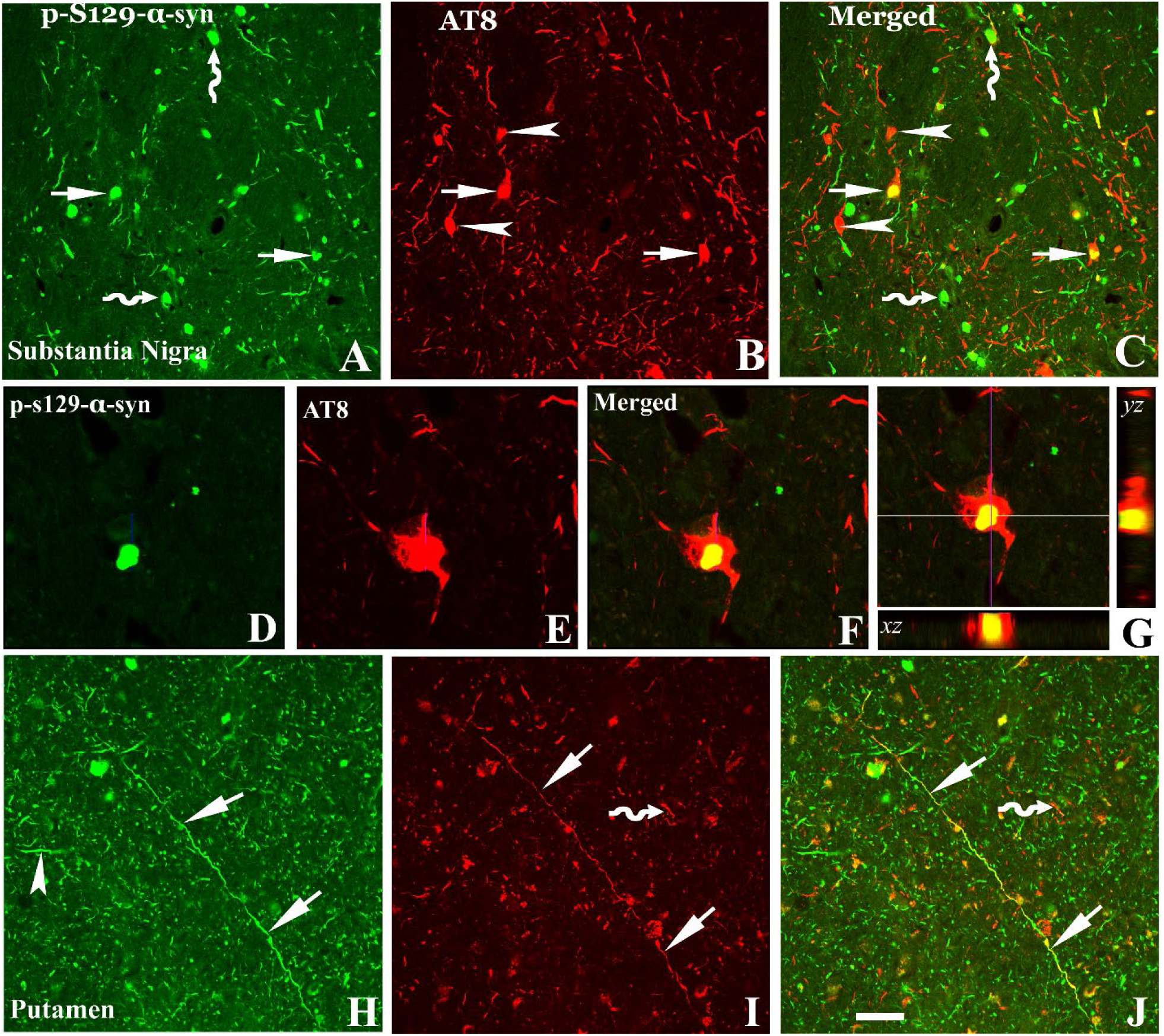
Co-localization analyses of phosphorylated tau (AT8) and phosphorylated α-synuclein (p-S129-α-syn) in nigrostriatal system. Confocal microscopic images of substantia nigra (A–G) and putamen (H–J) illustrated p-S129-α-syn (green; A, D, H), AT8 (red; B, E, I), and co-localization of AT8 and p-S129 (merged; C, F, G, J). There were three populations of immunofluorescence-labeling aggregates: p-S129-α-syn and AT8 double-labeling (arrows; A-C), AT8 single-labeling (arrowhead; B, C), or p-S129-α-syn single-labeling (curved arrow; A, C). The p-S129-α-syn-labeled aggregate (D) deposited within AT8-labeled perikarya (E, F). Three-dimensional reconstruction of confocal image further illustrated the colocalization of labeled p-S129-α-syn and AT8(G): the large panel represents a cross section of the cell layer, The horizontal (yellow) and vertical (pink) lines through them denote the planes of the adjoining xz and yz sections respectively. At the bottom and right, the xz and yz cross sections were obtained from the combined serial optical sections of these cell layers using Olympus Confocal Fluoroview software. The 3-dimensional reconstruction analyses revealed that p-S129-α-syn were colocalized with AT8 (yellow). In putamen p-S129-α-syn (arrowhead; H) and AT8-labeled threads and dots (curved arrow; I) were not colocalized but the longer fiber with AT8 labeling was p-S129-α-syn immunopositive (arrows; H-J). Scale bar = 50 μm in J (applies to H and I); 25μm for D-G; 100μm for A-C.

### Co-localization and quantitative analysis of dopaminergic markers in nigral neurons with tau aggregates

We know from our previous studies that experimental and human PD are associated with loss of dopaminergic phenotype in nigrostriatal neurons and believed at the time that this down-regulation was associated with α-syn [8, 9, 11]. Based upon the data described within, we now hypothesized that this down-regulation is tau dependent. To test this hypothesis, co-localization and quantitative analysis of TH immunoreactivity was performed in nigral neurons with or without tau aggregates from NMD, MMD, MMD-LB, and PD groups. Double-labelling revealed that the staining intensity of perikaryal TH immunoreactivity in nigral neurons without tau aggregates was similar across all four groups (Fig. 5). In contrast, nigral neurons with tau aggregates displayed reductions of TH immunoreactivity in subjects with MMD (Fig. 5D-5F), MMD-LB (5G-5I), and PD (Fig. 5J-5L). Quantitatively, an ANOVA revealed a significant difference in the optical density of TH immunoreactivity in nigral perikarya (p<0.0001) across groups. The fluorescence intensity of TH staining was reduced by 15.25-21.52% in neurons without tau aggregates in MMD, MMD-LB and PD groups and this was not statistically different from the NMD group (P>0.05). Interestingly in MMD, MMD-LB, and PD groups, nigral neurons that contained tau aggregates had significant reduction in TH immunofluorescence intensity (70.83% for MMD, 69.61% for MMD-LB, and 70.88% for PD) relative to NMD group (P<0.001; Fig. 5M).

**Figure 5.**
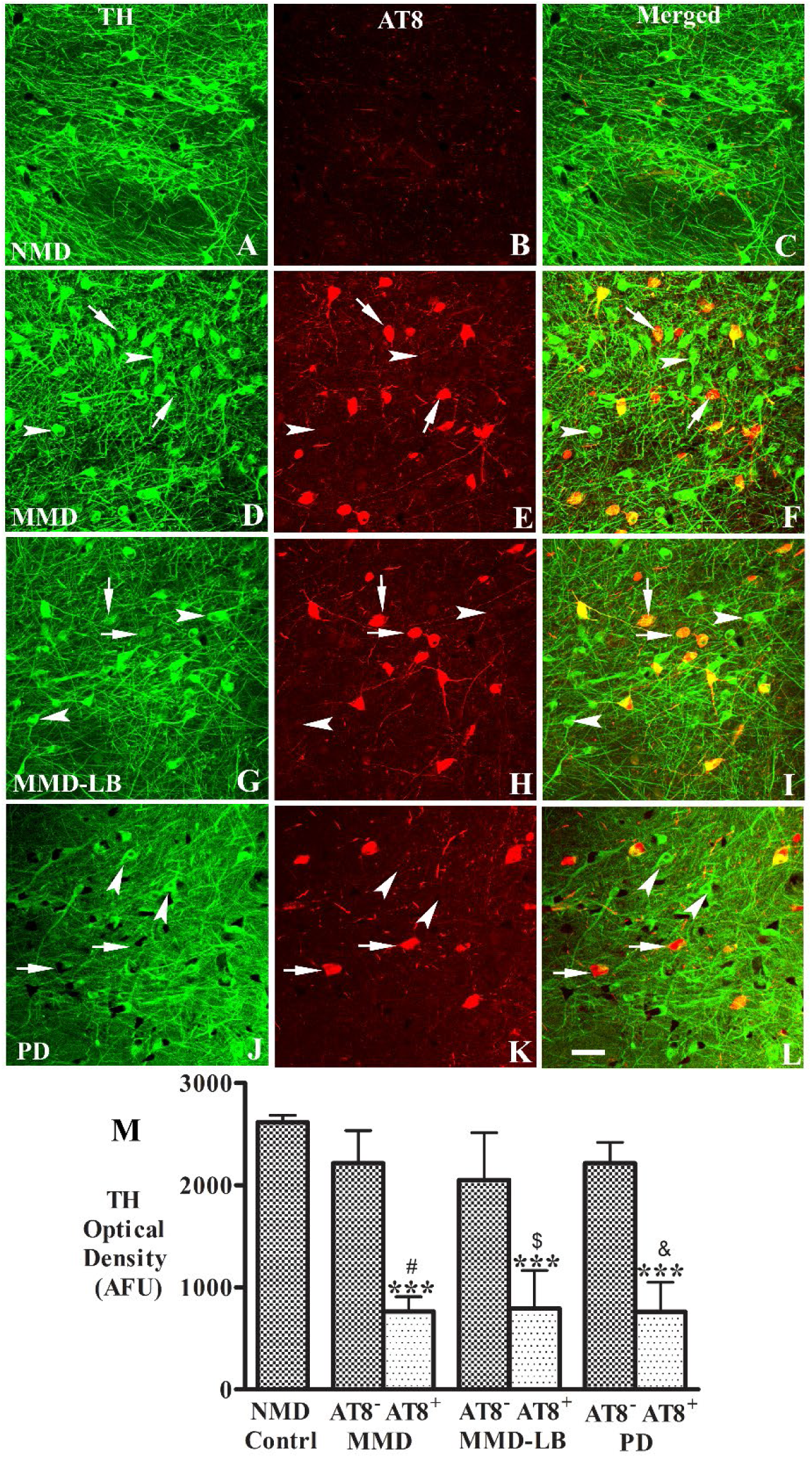
Reduction of tyrosine hydroxylase (TH) Levels in nigral neurons with AT8-ir aggregates. Confocal microscopic images of substantia nigra from no motor deficit (NMD; A-C), minimal motor deficits (MMD; D-F), minimal motor deficits with nigral Lewy body (MMD-LB; G-I) and Parkinson’s disease (PD); J-L) illustrated immunostaining for TH (green; A, D, G, J), AT8 (red; B, E, H, K), and co-localization TH and AT8 (merged; C, F, I, L). Note that TH immunofluorescent intensity was extensively reduced in the neurons with tau aggregates (arrows; D-L) but not in the neurons without tau aggregates (arrowheads; D-L). Scale bar in L = 100μm (applies to all). Measurements of immunofluorescent intensities (M) further revealed that TH expression was significantly reduced in the neurons with tau aggregates (AT8^+^) but not in the neurons without tau aggregate (AT8^-^). \*\*\**p*< 0.001 related to NMD control; ^#^*p*<0.05 related to AT8 immunonegative neurons in MMD, ^$^*P*<0.05 related to AT8 immunonegative neurons in MMD-LB, and ^&^*p*<0.05 related to AT8 immunonegative neurons in PD groups. Data are mean ± SD. AFU = arbitrary fluorescence units.

Within the putamen, a higher density of TH-labelled dopaminergic axons and terminals and no tau labelling were observed in NMD subjects (Fig. 6A-C). The TH-labeling was variable across the putamen in MMD (Fig. 6D) and MMD-LB (Fig. 6G). Some areas displayed TH-ir fibers with diminished intensity while others exhibited undetectable TH immunoreactivity as compared with NMD. PD (Fig. 6J) presented severe reductions of TH immunofluorescent intensity. The TH immunoreactivity was not detected in AT8-labled threads and puncta in all subjects with tau pathology (Fig. 6D-6L). Quantitative analyses revealed that the percent of TH-labeled products within the putamen was reduced 44.41% in MMD, 49.26% in MMD-LB, and 75.56%in PD group relative to NMD (Fig. 6M). An ANOVA revealed a statistically significant differences across these groups about intensity of a TH phenotype (P<0.0001). Post hoc analyses demonstrated that the optical density of TH-labeled fibers in the putamen was significantly reduced in MMD (P<0.001), MMD-LB (P<0.001) and PD (p<0.001) compared with NMD group, between MMD and PD (p<0.001), between MMD-LB and PD (P<0.001). In contrast, there was no significant difference between MMD and MMD-LB groups (P>0.05). These data indicate that tau, and not α-syn, drives the downregulation of TH in nigral neurons.

**Figure 6.**
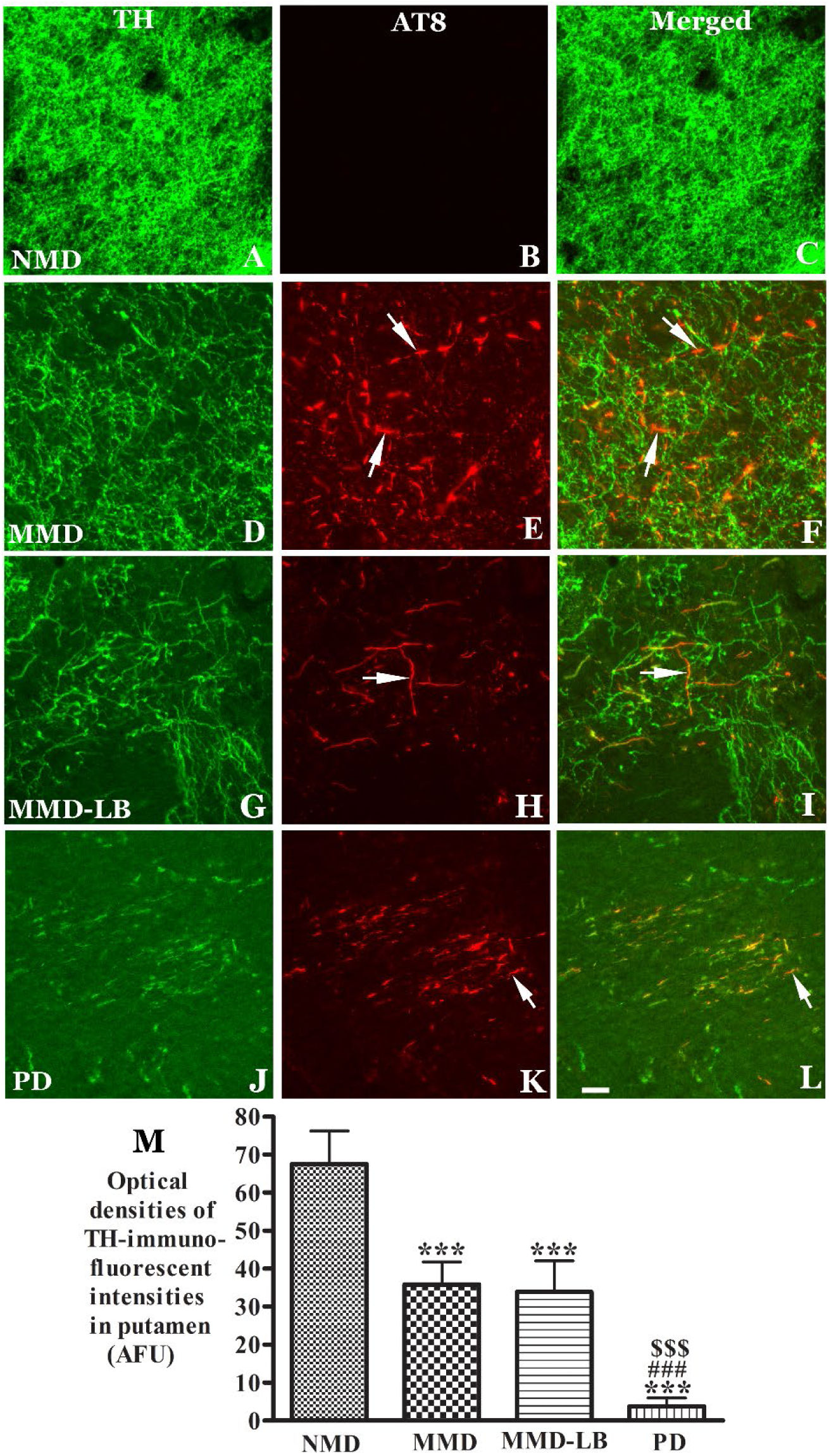
Reduction of tyrosine hydroxylase (TH) Levels in putamen with AT8-ir aggregates. Confocal microscopic images of putamen from no motor deficit (NMD; A-C), minimal motor deficits (MMD; D-F), minimal motor deficits with nigral Lewy body (MMD-LB; G-I) and Parkinson’s disease (PD; J-L) illustrated immunostaining for TH (green; A, D, G, J), AT8 (red; B, E, H, K), and co-localization TH and AT8 (merged; C, F, I, L). Note that TH immunofluorescent intensity was extensively reduced in MMD (D), MMD-LB (G) and PD groups (L) relative to NMD group (A). The AT8-labeled threads and punctuated dots (arrows, E, H, K) were TH immunonegative (arrows, F, I, L). Scale bar in L= 40μm (applies to all). Measurements of TH-ir intensity (M) further revealed that TH expression was significantly reduced in the putamen with tau aggregates in MMD, MMD-LB, and PD group. \*\*\**p*<0.001 related to NMD group; ^###^*p*<0.001 related to MMD group. ^$$$^*p*<0.001 related to MMD-LB group. Data are mean ± SD. AFU = arbitrary fluorescence units.

### Co-localization and quantitative analysis of kinesin light chain in nigral neurons with tau inclusion

Tau is active primarily in the distal portion of the axon where it provides microtubule stabilization and maintains routine axonal transport[29]. Normally tau is not detected in perikarya and dendrites although it is produced in perikarya[52]. In this study, tau inclusions existed in nigral perikarya and processes (Fig. 2, 3) in MMD, MMD-LB, and PD groups. However, phosphorylated tau inclusions were not detected in nigral perikarya and dendrites in NMD control group. The tau accumulation in nigral perikarya and dendrites represent misfolded tau protein. We have previously proposed that alterations in axonal transport is one of the earliest changes seen in nigrostriatal degeneration but had ascribed these changes as α-synuclein dependent[13]. Since the studies herein indicate that tau, and not necessarily α-synuclein, is correlated with nigrostriatal degeneration, we examined whether tau inclusions alter the expression of an axonal anterograde transport motor protein, kinesin light chain 1 (KLC) in the nigra of subjects in the NMD, MMD, and PD groups. Note that the MMD-LB group was not included in this analyses, as we have published reductions in KLC as a function of α-synuclein accumulation previously[13] and we did not have sufficient sections remaining in the MMD-LB group to perform this assessment rigorously. The subjects from MMD group were selected to analyze KLC alterations compared with subjects from NMD and PD groups. Double labelling with anti-tau (AT8) and anti-KLC antibodies (AP8637, ABGENT) was performed and optical density measurements for KLC were performed in neurons with or without tau inclusions. Co-localization studies revealed that both neurons with and without AT8-ir aggregates displayed significantly lower intensity of KLC immunoreactivity in MMD and PD relative to NMD (Fig. 7D). To further determine whether reductions in levels of KLC was associated with tau inclusions, we analyzed the relative intensity of KLC in nigral neurons that did or did not contain tau aggregates in the MMD group and compared these data to identically obtained data within NMD control and PD group. An ANOVA revealed a significant difference in optical density of KLC-ir intensity across these groups (Fig. 7J; p< 0.001). Post hoc analyses revealed a significant reduction of KLC-ir intensity in nigral neurons with (P<0.001) and without (P< 0.001) tau inclusions as compared with age-matched controls. However, the degree of KLC-ir was significantly (P<0.001) greater in neurons with tau inclusions relative to neurons without tau inclusions and these reductions were similar in MMD and PD groups (p>0.05).

**Figure 7.**
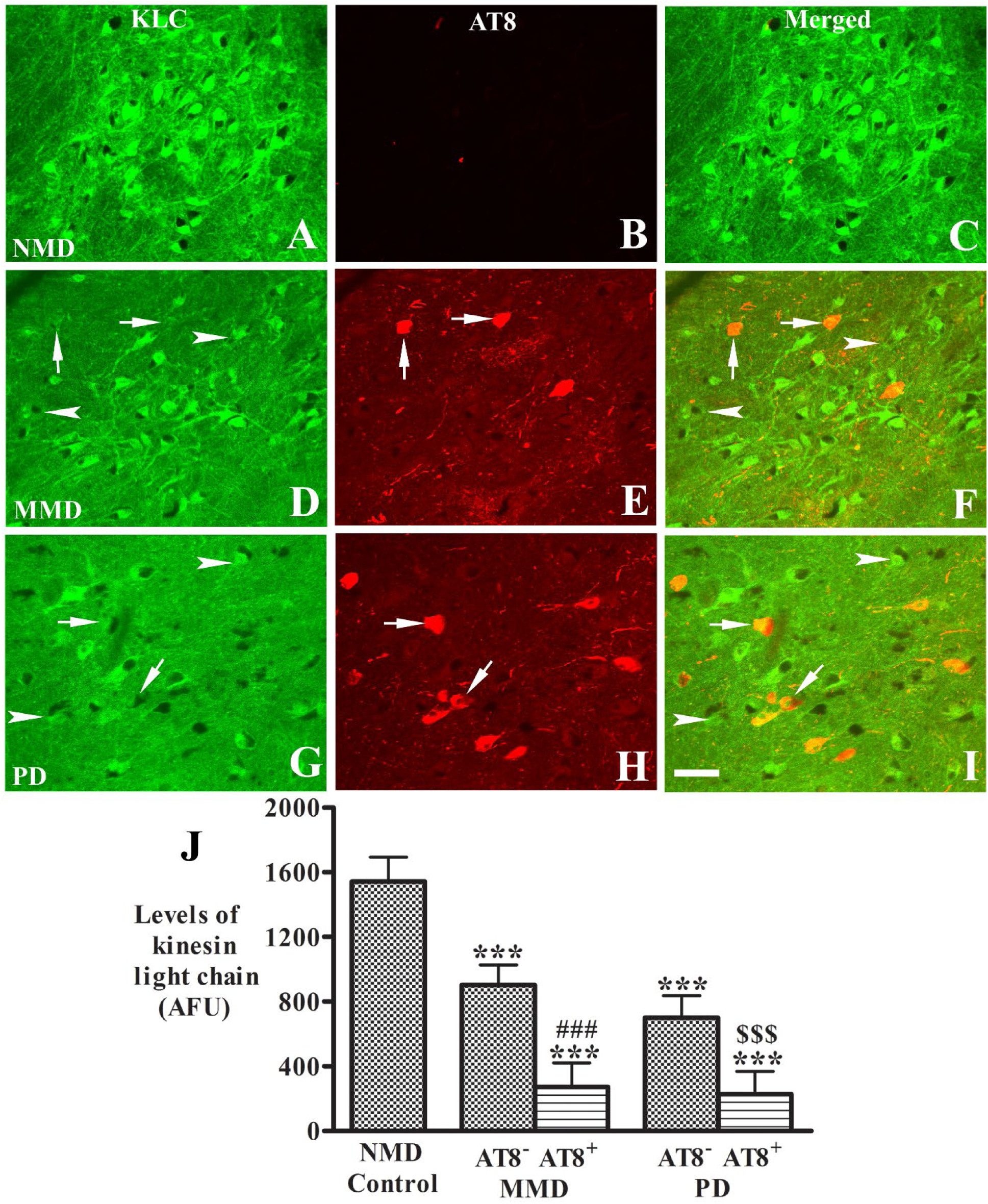
Reduction of kinesin light chain (KLC) levels in nigral neurons with tau aggregates. Confocal microscopic images of substantia nigra from no motor deficit (NMD; A-C), minimal motor deficits (MMD; D-F) and Parkinson’s disease (PD; H-I) illustrated immunostaining for KLC (green; A, D, G), AT8 (red; B, E, H), and co-localization KLC and AT8 (merged; C, F, I). Note that KLC immunofluorescent intensity was extensively reduced in both neurons with (arrows, D-1) and without tau aggregates (arrowheads; D-L). Scale bar in I = 100μm (applies to all). Measurements of immunofluorescent intensities (J) further revealed that KLC expression was significantly reduced in both neurons with (AT8^+^) and without tau aggregates (AT8^-^). \*\*\**p*<0.001related to NMD control group; ^###^*p*<0.001 related to AT8^-^ neurons in MMD group, ^$$$^*p*<0.01 related to AT8^-^ neurons in PD groups. Data are mean ± SD. AFU = arbitrary fluorescence units.

## Discussion

Lewy pathology and dopaminergic neuronal loss in the substantia nigra are defining pathologies in PD[5, 55]. However, several reports indicated that significant neurodegeneration and cellular dysfunction precede Lewy pathology emerging in the substantia nigra, suggesting these two events may not be pathologically linked[5, 7, 31, 40]. Further, there are emerging data from LRRK2 [30, 35] and other cases [58], that support the presence of clinical PD without synuclein pathology. Further, in support of this concept, subjects with Braak PD stages 1 and 2 exhibit dopaminergic neuronal dysfunction and neuronal loss, even though Lewy body pathology is undetectable in the substantia nigra[5, 40].

In the present study, we selected subjects with minimal parkinsonian signs, who were well characterized clinically as having parkinsonism but not PD and we view this population as prodromal PD. In all of our cases, a board-certified neuropathologist excluded other neurodegenerative diseases as contributory to our findings. In contrast to our previous studies, in which the cases were preselected based the clinical sign and the presence of Lewy pathology, this study included all, and then analyzed the status of the nigrostriatal system based on whether or not, they had Lewy pathology.

Seventeen cases with present nigral Lewy pathology were pathologically classified as MMD-LB.[17]. Eleven cases with parkinsonism, but without Lewy pathology, could be categorized as Braak PD stages 1 and 2 (MMD) because some had Lewy pathology in limbic and cortical areas, but not in the substantia nigra. Both MMD and MMD-LB groups with minimal clinical parkinsonian signs displayed intermediate reductions of TH-ir nigral neuron density and putamenal fibers relative to NMD controls and PDs. Critically, there was no statistically significant difference between subjects with clinical MMD regardless of whether they have Lewy pathology or not indicating that the clinical Parkinsonian syndrome and pathology occur independent of the Lewy pathology. Similarly, we investigated whether Lewy pathology was responsible for the downregulation of TH expression in substantia nigra perikarya and putamenal fibers during the development of PD. The present study also revealed that all brains with MMD cases had a decrease in their dopaminergic phenotype and these decreases in TH expression were identical in cases with and without Lewy pathology indicating further that Lewy pathology is not correlated with this form of degeneration.

What is the cellular insult causing nigral dopaminergic neuronal dysfunction and death that can precede Lewy body maturation? Several pathologies including tau, β-amyloid plagues, TDP-43, microinfarcts, atherosclerosis, arteriolosclerosis, and cerebral amyloid angiopathy have been associated with progression of parkinsonism in aging[7]. We hypothesized that phosphorylated tau accumulation appears to initiate nigral dopaminergic neurodegeneration as previous studies demonstrated neurofibrillary tangles in the substantia nigra in the elderly[50]. We observed pathological phospho-tau in all cases with MMD regardless of whether they had Lewy pathology or not. In fact, the degree of tauopathy was similar in MMD and MMD-LB subjects, suggesting that tau pathology, and not α-syn pathology, is the common denominator for what initiates the robust degeneration of the nigrostriatal pathway from the prodromal stage to PD. This is supported by the relatively lower densities of nigral tau-ir aggregates and putamenal tau-ir dots and threads in the PD group relative to the MMD and MMD-LB groups. At later stages of PD (H&Y5), tau-ir aggregates were rarely detected in nigrostriatal system. These nigrostriatal tau-ir aggregates in later stages of PD may disappear along with dopaminergic neuronal death. Tau aggregates accumulated in the nigrostriatal system in parkinsonism without nigral Lewy body pathology, suggesting that tau accumulation lies upstream of α-syn aggregates. Although the tau aggregates are not considered a prominent feature in PD pathogenesis[21], the prominent expression in this prodromal population warrants a re-examination of the sequencing of pathogenic events from preclinical, prodromal to manifest PD. Several studies reported that tau alone was sufficient to provoke severe neurodegeneration leading to parkinsonism in the absence of synucleinopathy in frontotemporal dementia and postencephalitic parkinsonism subjects[4, 34, 60]. Together these data support a mechanism where phosphorylated-tau accumulations are the initial pathogenic contributor for PD although the requirement of tau to lead to synucleinopathy will need to be established [54].

So what are the features of the tauopathy seen in this prodromal cohort of PD? There are multiple isoforms of tau with 3R-tau and 4R-tau being most prominent in humans. Both tau isoforms were seen in these cases and colocalized with phosphorylated tau marker AT8, indicating that the expression of 4R-tau was increased and subsequently phosphorylated. In the healthy brain, the 3R and 4R isoforms are equivalently expressed[26]. In Alzheimer’s brain, there is a decrease in 3R tau isoform or increase in 4R tau levels resulting in shift in the ratio of 4R-tau to 3R-tau[15, 25]. Whether the tau accumulation and aggregation in nigral dopaminergic neurons are associated with tau isoform imbalances needs further study.

Tau and α-syn are abundant brain proteins with distinct intraneuronal distribution and biological functions. Tau, a microtubule binding protein is restricted in axon where it stabilizes and promotes microtubule polymerization whereas α-syn is mainly localized in axon terminals where it may regulate synaptic functions. However, tau and α-syn are both partially unfolded proteins that can form toxic oligomers and abnormal intracellular aggregates under pathological conditions. The disorders with tauopathies are commonly accompanied with parkinsonian signs[1] while disorders with synucleinopathies are commonly attended by dementia[39]. Tau aggregates have been described in familial PD linked to A53T α-syn mutation[18, 36, 57] and in LRRK2 G2019S mutation carriers[30]. Our observations demonstrate that all MMD-LB cases have both pathologies as well as the vast majority of sporadic PD cases. Thus, while our MMD cases indicate that tau, and not α-syn, are critical for early manifestation of the disease, we cannot currently rule out and interaction in later disease. This morphological feature demonstrated that tau and α-syn may interact and the interaction may play an important role on the development and spreading of neurodegeneration[20, 28].

Mixed brain pathologies accelerate progression of PD development[7]. Cellular models have verified that pre-formed α-syn fibrils cross-seed intracellular tau to induce neurofibrillary tangle formation[23]. α-syn phosphorylation triggers tau pathogenicity and induces widespread phosphorylated tau with prion-like nature in various brain areas[28]. High levels of S396 phospho-tau and phospho-α-syn were found in synaptic-enriched fractions of the frontal cortex in Alzheimer’s disease and PD[42]. The SNCA gene polymorphism rs 2572324 has been reported relative to neocortical Lewy body and neurofibrillary tangle[48]. Our results support the concept that the α-syn and tau forms a deleterious feed-forward process essential for the development and spreading of neurodegeneration in PD and this occurs in a subset of subjects with prodromal PD. Genome-wide association study revealed that the genes encoding tau and α-syn have the highest association with PD[43, 44, 51]. Our results demonstrated that the majority of tau aggregates are not co-localized with α-syn aggregates and in fact a subpopulation of MMD subjects does not have nigral α-syn at al. Even if α-syn and tau aggregates occurred in the same brain, the formed aggregates were for the most part spatially separated. These data suggest that was no direct relationship between tau and α-syn aggregates in morphological analyses.

The misfolded protein accumulation and aggregation may be initially associated with other factors such as inefficiency of axonal transport. Tau is synthesized in nigral perikarya and transported to the distal portions of axons where it provides microtubule stabilization and flexibility as needed[14]. Tau is normally undetectable in perikarya and dendrites[29]. Our morphological analyses revealed phosphorylated tau accumulations from a few soluble granules to filling the nigral perikarya and processes with insoluble protein. Supported by these findings, we speculate that tau accumulation within perikarya, dendrites, and axons is associated with dysfunction of axonal transport. In our previous studies, in humans and PD models, we demonstrated decreases in the anterograde and retrograde axoplasmic transport motors, kinesin light chain and dynein respectively, are associated with α-syn aggregation. However, in the present study, we find decreases in kinesin light chain in neurons from MMD subjects with and without phosphorylated tau. Interestingly, we found a similar pattern of reduction in KLC in cases with PD, suggesting that other pathologies may impact this decrease but having misfolded tau or α-syn appears to exacerbate the effect[53].

In summary, the present series of observations is most consistent with pathological tau, and not pathological α-synuclein, initiating nigrostriatal degeneration in prodromal PD. Critically, this proposed mechanism hinges on MMD cases truly represent prodromal PD. The clinical and pathological data support this view. MMD cases are intermediate between NMD and PD in motor impairment, TH-ir nigral cell loss, loss of TH-ir putamenal innervation, nigral and putamenal phenotypic down-regulation. Alterations in kinesin-mediated impairment of axonal transport is the same in MMD and PD. All these pathological events are associated with tau pathology, and not α-syn, as they occur equally in MMD cases with and without Lewy pathology. These data should change our thinking of PD pathology and therapies directed towards reducing pathological tau might be a better strategy than those directed at α-syn. Furthermore, the use of an MMD cohort which displays robust clinical changes and pathological alteration on tau PET might be a cohort that would benefit most from such a clinical trial.

## Supporting information

https://www3.mydocsonline.com/Share.aspx?-379sxrPafWsSVyS3P8EI3UvuA

## Abbreviations

PD: Parkinson’s disease;
AT8: phosphor-Ser202+Thr205;
p-S129: phosphor-S129 α-synuclein;
TH: tyrosine hydroxylase;
NM: neuromelanin;
ir: immunoreactive;
α-syn: alpha-synuclein.

## Acknowledgments

We would like to thank Yinzhen He for histological assistance.

## Supplementary material

Supplementary material is available at *Acta Neuropathologica* online.

