## Supplementary material for "Is Tau the Initial Pathology in Dopaminergic Nigrostriatal Degeneration? Studies in Parkinsonism and Parkinson’s Disease": https://www3.mydocsonline.com/Share.aspx?-379sxrPafWsSVyS3P8EI3UvuA

### Supplementary Figures

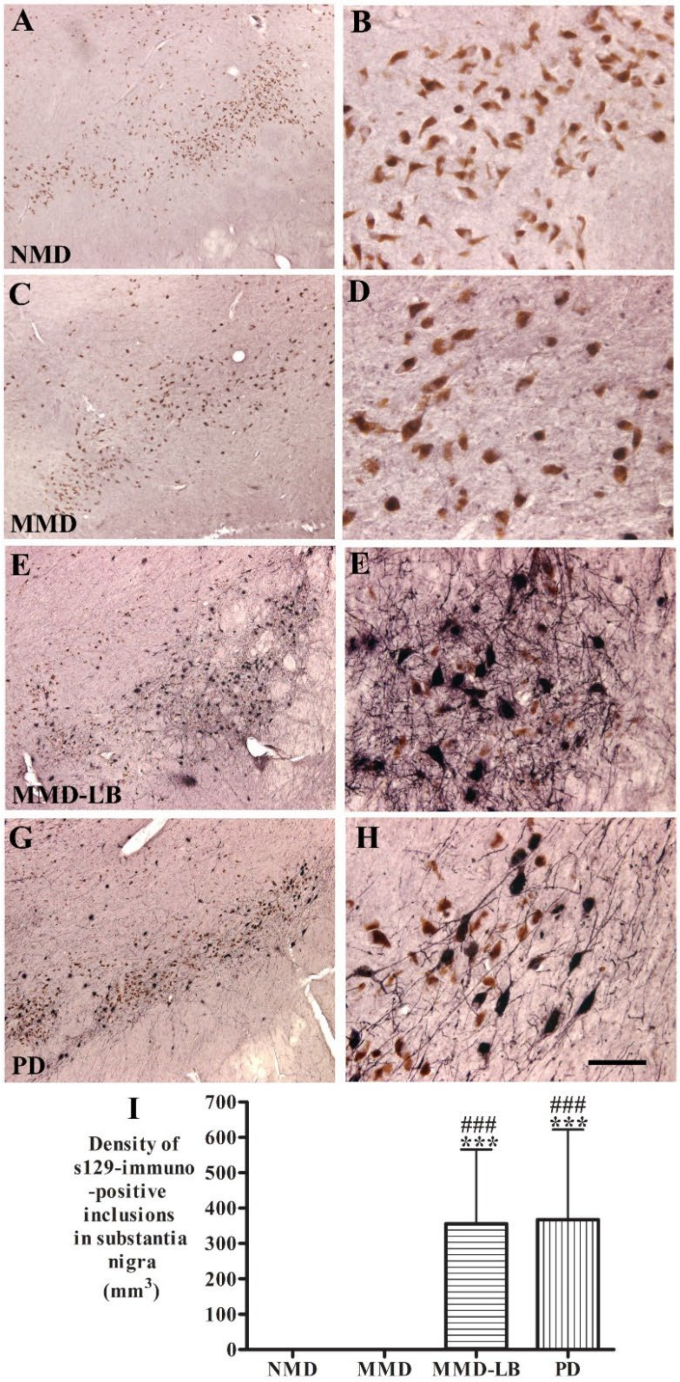


**Supplementary Figure 1. Qualitative and quantitative evaluation for phosphorylated alpha-synuclein aggregates in substantia nigra.**

Photomicrographs of the mid-substantia nigra from no motor deficit (NMD; A, B), minimal motor deficits (MMD; C, D), minimal motor deficits with nigral Lewy body (MMD-LB; E, F), and Parkinson's disease (PD; G, H) show phosphor-S129 α-synuclein (p-S129) patterns. p-S129 immunoreactivity was undetectable in the subjects with NMD (A, B) and MMD (C, D). In contrast, p-S129-immunoreactive nigral neurons were observed in subjects with MMD-LB (E, F) and patients with PD (G, H). Note there were more p-S129-a-syn–immunoreactive processes in subjects with MMD-LB (E) than in patients with PD (H). Scale bar in H = 100 μm for B, D, F; 500 μm for A, C, E, G. (I) Stereological analyses revealed that no p-S129-positive aggregate was counted in NMD and MMD groups. However, there was a significantly higher densities of p-S129 labeled aggregates in MMD-LB and PD groups. *** $p<0.001$ compared with NMD; $^{###}p<0.001$ compared with MMD.

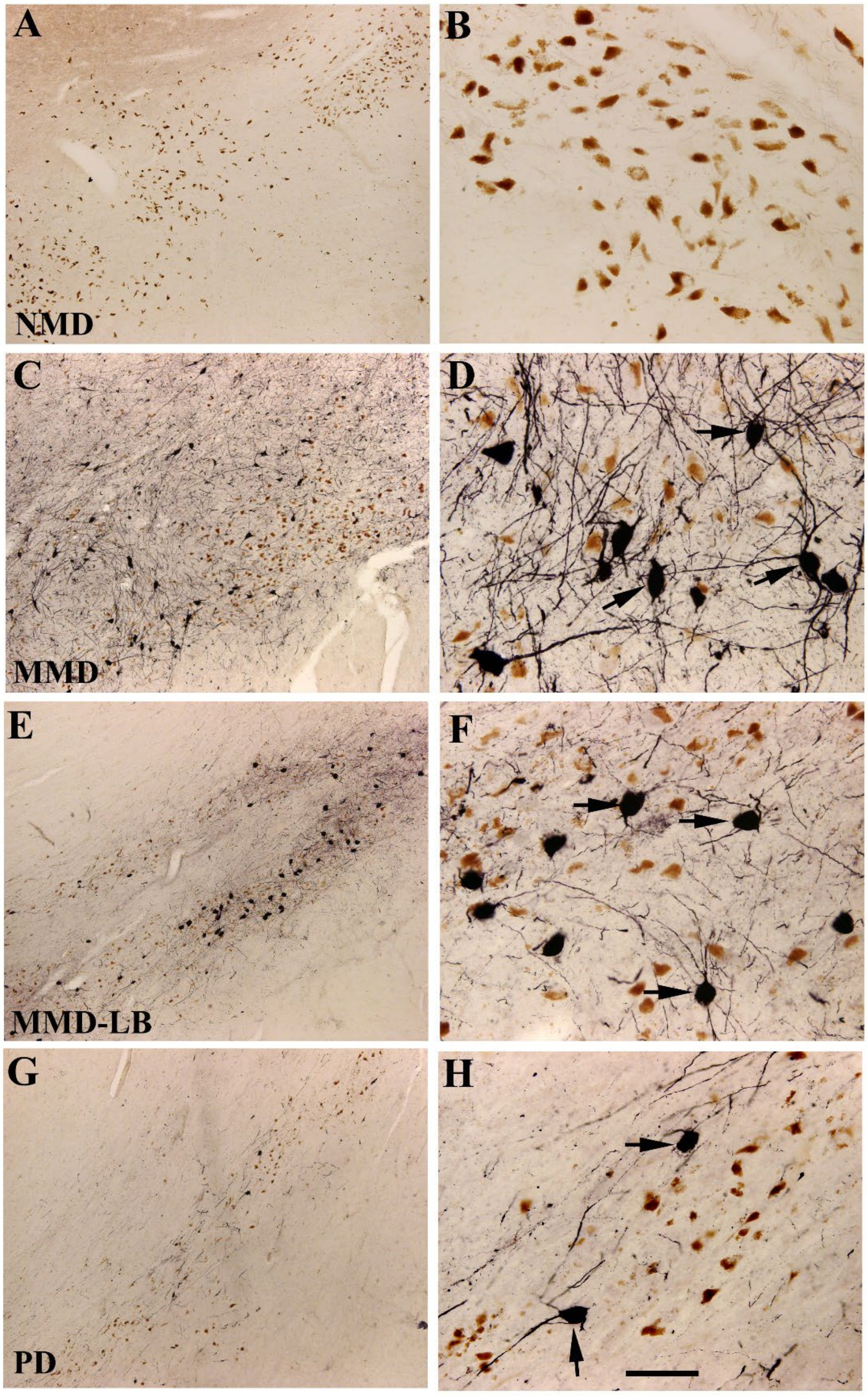


**Supplementary Figure 2. Qualitative observation for tau phosphorylated at serine-202 in substantia nigra.** Photomicrographs of the mid-substantia nigra from no motor deficit (NMD; A, B), minimal motor deficits (MMD; C, D), minimal motor deficits with nigral Lewy body (MMD-LB; E, F), and Parkinson's disease (PD; G, H) show phosphorylated Serine-202 (CP13) tau patterns. CP13 immunoreactivity was undetectable in the subjects with NMD (A, B). In contrast, CP13-immunoreactive nigral neurons were observed in subjects with MMD (C, D), MMD-LB (E, F), and patients with PD (G, H). Note there were more CP13–immunoreactive neurons with extensive processes in subjects with MMD (arrows, D) and MMD-LB (arrows, F) than the patients with PD (arrows, H). Scale bar in H = 100 µm for B, D, F; 500 µm for A, C, E, G.

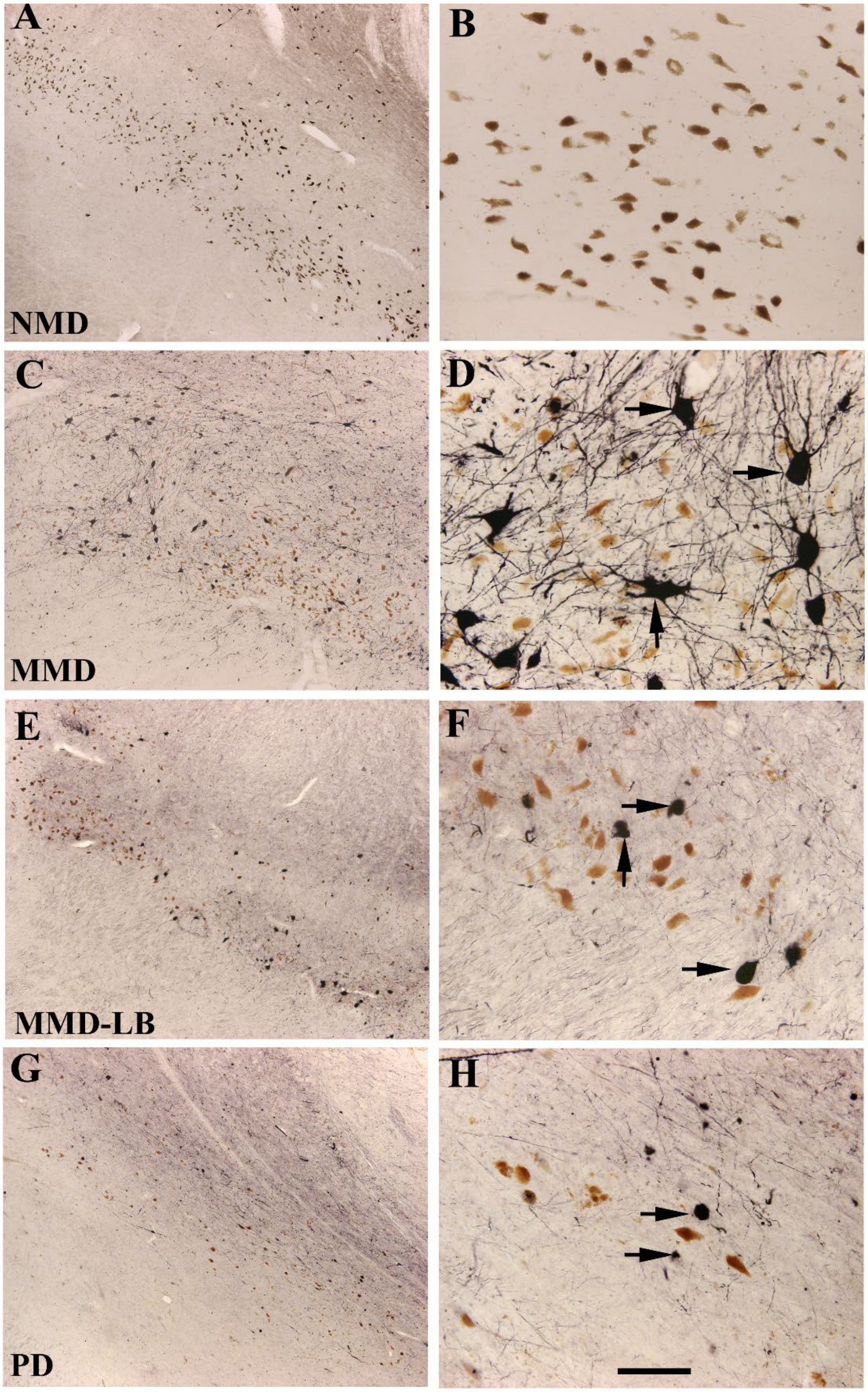


**Supplementary Figure 3. Qualitative observation for tau phosphorylated at Ser396/Ser404 in substantia nigra.** Photomicrographs of the mid-substantia nigra from no motor deficit (NMD; A, B), minimal motor deficits (MMD; C, D), minimal motor deficits with nigral Lewy body (MMD-LB; E, F), and Parkinson's disease (PD; G, H) show phosphorylated Ser396/Ser404 (PHF-1) tau patterns. PHF-1 immunoreactivity was detected in the subjects with MMD (C, D), MMD-LB (E, F) and PD (G, H) but not NMD (A, B). Note that PFH-1–immunoreactive perikarya with extensive processes in subjects with MMD (arrows, D) while restricted processes in MMD-LB (arrows, F) and PD (arrows, H). Scale bar in H = 100 µm for B, D, F; 500 µm for A, C, E, G.

### Supplementary Tables

#### Supplementary Table 1. Demographics of older adults with no motor deficit (NMD)

| Case no. | Age (years) | Gender | PMI (hour) | Glob-P (0-100) | Gait (0-100) | Rigidity (0-100) | Tremor (0-100) | Bradykinesia (0-100) |
|---|---|---|---|---|---|---|---|---|
| B00-76 | 71 | M | 4.08 | 6.51 | 0 | 26.06 | 0 | 0 |
| B97-32 | 73 | M | 7.25 | 3.47 | 0 | 0 | 13.91 | 0 |
| B98-115 | 84 | M | 4.00 | 2.40 | 9.63 | 0 | 0 | 0 |
| B96-98 | 72 | M | 7.00 | 3.03 | 12.14 | 0 | 0 | 0 |
| B98-84 | 78 | F | 3.00 | 2.58 | 0 | 10.35 | 0 | 0 |
| B00-59 | 91 | F | 10.66 | 4.90 | 0 | 0 | 0 | 19.66 |
| B99-07 | 84 | F | 3.50 | 2.14 | 8.57 | 0 | 0 | 0 |
| B96-50 | 88 | F | 7.00 | 6.35 | 25.42 | 0 | 0 | 0 |
| AT7-1 | 91 | F | 6.50 | 7.58 | 30.32 | 0 | 0 | 0 |
| **Mean±SD** | **81.33±8.06** | **4/5** | **5.88±2.45** | **4.32±2.05** | **9.56±11.48** | **4.04±8.93** | **1.54±4.63** | **2.18±6.55** |

PMI, postmortem interval; MMSE, Glo-P, global parkinsonism.

#### Supplementary Table 2. Demographics of older adults with motor deficit but absent nigral Lewy body (MMD)

| Case no. | Age (years) | Gender | PMI (hour) | Glob-P (0-100) | Gait (0-100) | Rigidity (0-100) | Tremor (0-100) | Bradykinesia (0-100) |
|---|---|---|---|---|---|---|---|---|
| AT2-5 | 95 | F | 6.17 | 15.80 | 35.71 | 0.00 | 0.00 | 27.50 |
| AT3-1 | 97 | F | 10.08 | 21.31 | 66.67 | 0.00 | 6.06 | 12.50 |
| AT4-1 | 87 | F | 4.92 | 12.5 | 25.00 | 25.00 | 0.00 | 12.50 |
| AT5-1 | 96 | F | 12.61 | 32.41 | 82.14 | 20.00 | 0.00 | 27.50 |
| AT5-3 | 92 | M | 14.17 | 29.23 | 64.29 | 10.00 | 15.15 | 27.50 |
| AT5-4 | 92 | F | 7.41 | 9.29 | 32.14 | 0.00 | 0.00 | 5.00 |
| AT6-1 | 90 | F | 18.75 | 34.82 | 64.29 | 30.00 | 0.00 | 45.00 |
| AT6-3 | 95 | F | 7.1 | 44.01 | 78.57 | 60.00 | 0.00 | 37.50 |
| AT6-4 | 93 | M | 12.75 | 23.84 | 64.29 | 0.00 | 6.06 | 25.00 |
| AT6-5 | 96 | F | 6.5 | 41.60 | 71.43 | 35.00 | 0.00 | 60.00 |
| AT6-6 | 90 | M | 9.75 | 14.28 | 42.11 | 0.00 | 0.00 | 15.00 |
| **Mean±SD** | **93.00±3.10** | **3/8** | **10.24 ±4.07** | **25.37 ±11.92** | **56.96 ±19.68** | **16.36±19.76** | **2.47±4.85** | **26.81±16.05** |

PMI, postmortem interval; Glo-P, global parkinsonism.

**Supplementary Table 3. Demographics of older adults with minimal motor deficit and nigral Lewy body (MMD-LB)**

| Case no. | Age (years) | Gender | PMI (hour) | Glob-P (0-100) | Gait (0-100) | Rigidity (0-100) | Tremor (0-100) | Bradykinesia (0-100) |
|---|---|---|---|---|---|---|---|---|
| AT1-1 | 92 | F | 4.75 | 50.00 | 57.14 | 80.00 | 20.00 | 17.50 |
| AT1-2 | 88 | F | 8.17 | 30.00 | 42.86 | 60.00 | 0.00 | 2.50 |
| AT1-3 | 89 | M | 0.75 | 19.82 | 39.29 | 20.00 | 0.00 | 20.00 |
| AT2-1 | 95 | F | 26.05 | 20.71 | 0.00 | 15.00 | 12.12 | 35.00 |
| AT2-2 | 95 | M | 5.67 | 13.75 | 50.00 | 5.00 | 0.00 | 0.00 |
| AT2-3 | 90 | F | 6.67 | 8.40 | 26.09 | 0.00 | 0.00 | 7.50 |
| AT2-4 | 97 | M | 4.75 | 15.95 | 57.14 | 25.00 | 0.00 | 22.86 |
| AT2-6 | 100 | F | 6.33 | 16.19 | 75.00 | 10.00 | 6.06 | 32.50 |
| AT2-7 | 85 | M | 6.50 | 9.91 | 32.14 | 5.00 | 0.00 | 2.50 |
| AT3-2 | 89 | M | 5.33 | 8.70 | 14.29 | 5.00 | 3.03 | 12.50 |
| AT3-3 | 90 | M | 13.92 | 37.85 | 67.86 | 30.00 | 6.06 | 47.50 |
| AT3-4 | 94 | F | 4.40 | 10.87 | 43.48 | 0.00 | 0.00 | 0.00 |
| AT3-5 | 93 | F | 13.52 | 5.09 | 17.86 | 0.00 | 0.00 | 2.50 |
| AT3-6 | 90 | M | 5.58 | 10.35 | 38.13 | 15.00 | 6.06 | 10.00 |
| AT4-2 | 82 | M | 17.00 | 8.21 | 17.86 | 0.00 | 6.06 | 15.00 |
| AT4-3 | 77 | M | 13.90 | 8.75 | 0.00 | 0.00 | 0.00 | 35.00 |
| AT4-4 | 91 | F | 16.00 | 3.39 | 3.57 | 5.00 | 0.00 | 5.00 |
| **Mean±SD** | **90.41±5.53** | **9/8** | **9.37±6.37** | **16.34±12.41** | **34.27±23.15** | **16.18±22.54** | **3.49±5.56** | **15.75±14.47** |

PMI, postmortem interval; Glo-P, global parkinsonism.

Supplementary Table 4. Demographics of Patients with Parkinson's Disease

| Case. No | Age (years) | Gender | PMI (hour) | UPDRSIII (on) | H&Y (on) | Glob-P (0-100) | Gait (0-100) | Rigidity (0-100) | Tremor (0-100) | Bra-K (0-100) |
|---|---|---|---|---|---|---|---|---|---|---|
| B13-45 | 57 | M | 6.30 | 35.0 | 2 | 25.17 | 25.00 | 15.00 | 10.71 | 50.00 |
| B12-95 | 66 | M | 12.00 | 38.0 | 2 | 32.12 | 20.00 | 25.00 | 42.86 | 40.63 |
| B10-90 | 77 | F | 3.50 | 22.0 | 2 | 24.93 | 25.00 | 10.00 | 17.86 | 46.88 |
| B09-04 | 74 | M | 8.21 | 38.5 | 2 | 25.47 | 50.00 | 30.00 | 0.00 | 21.88 |
| B14-01 | 71 | M | 5.30 | 33.5 | 3 | 27.32 | 37.50 | 20.00 | 14.28 | 37.50 |
| B13-46 | 70 | F | 6.00 | 51.0 | 3 | 46.63 | 60.00 | 60.00 | 7.14 | 59.38 |
| B12-66 | 87 | F | 6.00 | 31.0 | 3 | 27.14 | 30.00 | 25.00 | 28.57 | 25.00 |
| B10-10 | 77 | F | 5.00 | 53.0 | 3 | 41.69 | 55.00 | 35.00 | 14.29 | 62.50 |
| B12-52 | 89 | M | 2.30 | 39.0 | 3 | 31.87 | 40.00 | 50.00 | 0.00 | 37.50 |
| B13-49 | 80 | M | 10.00 | 46.0 | 4 | 44.06 | 70.00 | 50.00 | 0.00 | 56.25 |
| B13-47 | 86 | M | 6.30 | 53.0 | 4 | 39.37 | 50.00 | 45.00 | 0.00 | 62.50 |
| B13-50 | 72 | M | 7.00 | 54.0 | 5 | 47.18 | 95.00 | 50.00 | 0.00 | 43.75 |
| B04-42 | 76 | M | 3.30 | 65.0 | 4 | 47.18 | 70.00 | 50.00 | 0.00 | 68.75 |
| B07-11 | 72 | F | 5.00 | 58.0 | 5 | 54.34 | 66.66 | 65.00 | 0.00 | 85.71 |
| B08-110 | 95 | F | 3.30 | 49.0 | 5 | 45.11 | 83.33 | 40.00 | 0.00 | 57.14 |
| B09-127 | 76 | F | 4.15 | 44.0 | 4 | 46.09 | 75.00 | 50.00 | 0.00 | 59.38 |
| B12-110 | 88 | F | 5.00 | 42.0 | 4 | 38.75 | 55.00 | 50.00 | 0.00 | 50.00 |
| B14-04 | 81 | M | 7.30 | 40.0 | 4 | 33.75 | 50.00 | 35.00 | 0.00 | 50.00 |
| B14-05 | 85 | F | 4.30 | 37.0 | 3 | 26.67 | 55.00 | 20.00 | 3.57 | 28.13 |
| B16-02 | 74 | F | 10.00 | 40.0 | 4 | 28.56 | 16.66 | 15.00 | 35.71 | 46.87 |
| B95-129 | 70 | M | 9.00 | 47.0 | 4 | 41.75 | 79.16 | 20.00 | 0 | 67.85 |
| B96-11 | 77 | M | 5.30 | 42.0 | 3 | 43.75 | 75.00 | 50.00 | 0 | 50.00 |
| B10-02 | 86 | M | 4.00 | 65 | 5 | 62.57 | 83.33 | 75.00 | 6.25 | 85.71 |
| B98-09 | 87 | F | 9.00 | 62.0 | 5 | 45.95 | 79.16 | 35.00 | 12.50 | 57.14 |
| **Mean±SD** | **78.04±8.69** | **13/11** | **5.99±2.41** | **45.20±10.91** | **3.58±1.01** | **38.64±10.24** | **56.07±22.34** | **38.33±17.29** | **8.07±12.27** | **52.10±16.28** |

PMI, postmortem interval; UPDRS, United Parkinson's Disease Rating Scale; Bra-k, Bradykinesia
